# Single-molecule motion profiling reveals direct and indirect effects of PolyQ expansion on huntingtin dynamics and molecular engagement

**DOI:** 10.64898/2026.11.25.769260

**Authors:** Michael Shannon, Shu Li, Thi Ngo, Tiago L. Laundos, Catherine Kim, Francesco M. Piccolo, Ali H. Brivanlou

## Abstract

Huntington’s disease (HD) is caused by CAG expansion in *HTT*, but how PolyQ expansion alters huntingtin behaviour in human neurons remains unclear. We combined live single-molecule imaging and tracking with transformer-based motion profiling to resolve full-length HTT dynamics in pluripotent hESCs and their derived cortical- and MSN-like neurons. HTT occupied three behavioural states: bound, confined, and exploratory. PolyQ expansion progressively shifted molecules from bound and confined states toward the exploratory state. These effects emerged at shorter PolyQ lengths in MSN-like neurons, and only at 150Q included faster exploratory diffusion and impaired return to stable bound or confined states. Tag-swap experiments showed that mt-HTT imposed similar defects on co-expressed wild-type HTT (wt-HTT), revealing a dominant effect of the HD mutation on wt-HTT dynamics. Loss of HTT (HTT-/-) or expression of mt-HTT also altered kinesin and myosin dynamics, consistent with changes in the intracellular environment of HD cells. Thus, PolyQ-expanded HTT does not merely acquire abnormal dynamics, it leads to a broader disruption of the cytoplasmic environment in which both wt-HTT and molecular motors operate.

## Introduction

Huntington’s disease (HD) is caused by expansion of CAG repeats in exon 1 of the huntingtin *(HTT)* gene, producing an expanded polyglutamine (PolyQ) tract (MacDonald et al. 1993). PolyQ lengths above 35CAG repeats are associated with disease, with increasing repeat length linked to earlier clinical onset (Kay et al. 2016; Andrew et al. 1993; M. Duyao et al. 1993). Although multiple brain regions are affected, striatal medium spiny neurons (MSNs) show early and pronounced vulnerability, followed by broader cortical degeneration (Reiner et al. 1988; Aylward, Nopoulos, Ross, Langbehn, Pierson, Mills, Johnson, Magnotta, Juhl, Paulsen, and PREDICT-HD Investigators and Coordinators of Huntington Study Group 2011). How PolyQ expansion alters normal HTT function to produce this selective vulnerability remains incompletely understood.

Wildtype HTT (wt-HTT) is a large, ubiquitously expressed protein required for development and neuronal function. Complete loss is embryonic lethal in mice (M. P. Duyao et al. 1995), whereas depletion in the adult nervous system causes progressive neurodegeneration (Dragatsis et al. 2000). wt-HTT has been implicated in trafficking, survival, polarity, ciliogenesis and cytoskeletal organization, consistent with activity mediated through diverse molecular partners and intracellular structures (Caviston et al. 2007; Zhang et al. 2006; Galgoczi et al. 2021; Saudou and Humbert 2016; Gauthier et al. 2004; Haremaki et al. 2015; Molina-Calavita et al. 2014; Capizzi et al. 2022; Carpentier et al. 2025). Accordingly, wt-HTT function may depend not only on which partners it encounters, but on where, when and for how long those interactions are maintained. Molecular motion should therefore contain information about these engagements. Thus, wt-HTT molecular dynamics can serve as a functional readout of its engagement with the intracellular environment.

In our previous studies we generated a collection of isogenic human embryonic stem cell lines (hESCs) carrying different CAG-repeat lengths (mt-HTT), as well as a null (HTT-/-) line. Embryo models and neuronal organoids derived from these lines revealed PolyQ-length dependent phenotypic signatures, which phenocopied those of HTT-/-, suggesting that HD-mutation represents a loss-of-function (Haremaki et al. 2019; Galgoczi et al. 2021; Piccolo et al. 2022). More recently, we showed that mt-HTT can interfere with the activity of co-expressed wt-HTT, supporting a dominant component of the HD mutation (Laundos et al. 2023). Distinguishing further these defects intrinsic to mt-HTT from effects imposed on wt-HTT therefore requires observing the two proteins independently, within the same cell. Their molecular dynamics provide a means to test whether PolyQ expansion changes HTT because the mutant protein itself behaves differently, or because it changes the environment in which HTT moves/interacts, or both.

Single-particle tracking can resolve nanoscale dynamics in living cells, but molecular trajectories are intrinsically heterogeneous. Conventional quantities such as the diffusion coefficient, speed and turning angle collapse complex trajectories into scalar measurements, obscuring the behaviours that generate them (Kowalek et al. 2019). An increase in average diffusion, for example, could reflect faster movement of individual molecules, a reduction in the stably bound population, changes in switching between bound and mobile states, or some combination of these effects. Feature based classification can separate some heterogeneity but requires preselected parameters and imposes discrete boundaries between behavioural classes (Pinholt et al. 2021). Because individual molecules can change behaviour over time and occupy intermediate regimes (Vega et al. 2018), an analysis framework should retain both the sequence of motions and uncertainty in behavioural assignment inherent to this continuously labile system.Transformers provide one approach because they learn contextual relationships within ordered sequences (Vaswani et al. 2017). Molecular trajectories can similarly be treated as sequences in which successive displacements and directional changes acquire meaning from the motions that precede and follow them, analogous to words acquiring meaning from their context within a sentence. This allows behaviour to be learned directly from trajectory structure rather than solely from hand selected scalar features.

Here, using wt-and mt-HTT with kinesin and myosin as reference proteins, we define four interpretable motion states: bound, confined, exploratory and transport-like, and train a raw-motion transformer to classify these states directly from local trajectory sequences. Applying this framework to hESCs and hESC-derived cortical and MSN-like neurons allow us to examine how HTT dynamics vary across cellular and neuronal compartments and whether disease-relevant neuronal identities differ in their response to PolyQ expansion.

Monitoring the live activity of differentially tagged full-length HTT proteins, we distinguish changes intrinsic to mt-HTT from those imposed on co-expressed wt-HTT. Namely, increasing PolyQ length progressively shifts mt-HTT away from bound and confined states toward exploratory motion, with effects emerging at shorter PolyQ lengths in MSN-like compared with cortical neurons. At 150Q and not below, a fast exploratory population becomes more mobile and HTT shows altered motion-state switching over longer timescales. Interestingly, mt-HTT imposes related behavioural defects on co-expressed wt-HTT, extending the dominant effect previously described in stem-cell patterning and neuruloid models (Laundos et al. 2023) to single-molecule dynamics.

Extending the analysis to kinesin and myosin shows that mt-HTT and HTT-/- alters the dynamics of molecular motor proteins, supporting broader disruption of the cytoplasmic environment. Together, these findings connect PolyQ expansion to both intrinsic disruption of HTT molecular engagement and indirect remodelling of the environment in which HTT and other proteins operate.

## Results

### Comparative baseline motion of HTT in pluripotent hESCs

To compare wt-HTT and mt-HTT dynamics at single-molecule resolution, we used full-length HTT carrying a C-terminal HaloTag. We previously demonstrated that C-terminally tagged wt-HTT retains functional activity, rescuing HTT-knockout phenotypes in micropatterned and neuruloid assays (Laundos et al. 2023). To establish a baseline for interpreting the dynamic features of HTT proteins we compared the dynamics of exogenously expressed wt-HTT-HaloTag in the HTT-/- background with that of endogenously tagged HTT in the undifferentiated pluripotent wt-human embryonic stem cells(hESCs; RUES2-NIHhESC-09-0013). None of the measured parameters: diffusion coefficient, speed, self-intersections, cumulative displacement or anomalous exponent were significantly different between the endogenously tagged and exogenously expressed wt-HTT (Figure. S1), indicating that under our experimental conditions, exogenous expression does not measurably alter the baseline nanoscale dynamic behaviour of HTT. We then benchmarked wt-HTT motion against two well-characterized cytoskeletal motor proteins: Halo-tagged KIF5B (kinesin) and MYH9 (myosin) in living hESCs); using lines selected for homogenous expression (Figure S2). Single molecules were imaged at 100 Hz (Figure 1A) and trajectories segmented into equal 600-ms windows for comparison (Figure 1B). All three proteins displayed heterogeneous motion, including highly localized, confined and broadly exploratory trajectories, while processive directional motion was most evident for kinesin (Figure 1B).

**Figure 1.**
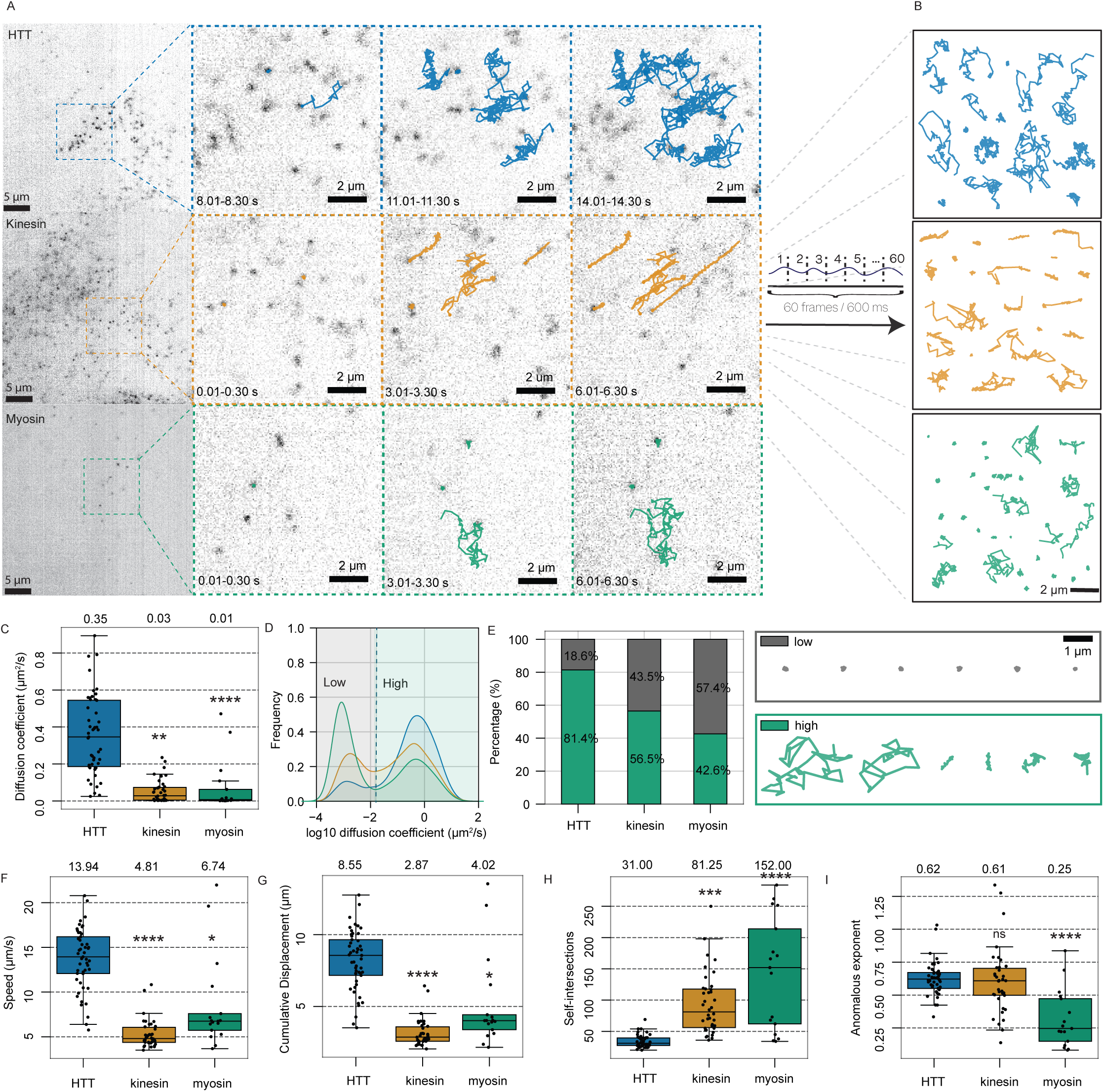
Single HTT molecules move rapidly and predominantly exhibit exploratory motion compared with kinesin and myosin. (A) Halo-tagged HTT (blue), kinesin (orange) and myosin (green) imaged in undifferentiated human embryonic stem cells (hESCs) by TIRF microscopy. Enlarged regions show subsets of single-molecule trajectories recorded over 6 s from 60-s movies for each protein. (B) Representative trajectories of HTT, kinesin and myosin displayed over the same interval (60 frames; 600 ms) and separated for clarity. (C) Diffusion coefficients for each protein; dots represent cell-level medians. (D) Histograms of log10-transformed diffusion coefficients for HTT, kinesin and myosin, with the threshold separating low- and high-diffusion populations. (E) Percentages of trajectory windows in the low- and high-diffusion populations for each protein. (F–I) Boxplots of additional trajectory features: (F) speed, (G) cumulative displacement, (H) number of self-intersections and (I) anomalous exponent. Dots represent cell-level medians. Sample sizes: HTT, 36 cells and 29,196 windowed trajectories; kinesin, 12 cells and 10,808 windowed trajectories; myosin, 12 cells and 6,163 windowed trajectories. Combined total: 60 cells, 12,328 tracks, 46,167 windows and 1,708,920 data points. Statistical significance was assessed using the Kruskal–Wallis test followed by Dunn’s post hoc test with Bonferroni correction. **p < 0.001, ***p < 0.0001, ****p < 0.00001; ns, not significant.

wt-HTT was substantially more mobile than kinesin or myosin in undifferentiated pluripotent hESCs, with a median diffusion coefficient of 0.35 um^2^/s compared with 0.03 and 0.01 um^2^/s respectively (Figure 1C). Log10 diffusion coefficient distributions were bimodal for all three proteins but differed markedly in composition. Using the minimum between the two peaks to define the boundary between low- and high-diffusion populations (0.016 µm²/s), 81% of HTT trajectories belonged to the high-diffusion population, compared with 56% of kinesin and 42% of myosin (Figure 1D, E). The low diffusion population consisted predominantly of highly localized trajectories, whereas the higher diffusion population contained more heterogeneous motion (Figure 1E). Consistently, wt-HTT moved faster (Figure 1F), travelled farther (Figure 1G) and showed fewer self-intersections (Figure 1H) than kinesin or myosin. wt-HTT and kinesin also showed higher anomalous exponents (α), indicating less constrained motion approaching Brownian diffusion, whereas myosin showed lower α values indicative of more constrained, sub-diffusive behaviour (Figure 1I).

Together, wt-HTT, kinesin and myosin sampled the same broad repertoire of molecular motion but with distinct dynamic profiles. wt-HTT was dominated by mobile, exploratory trajectories, myosin by highly localized motion, and kinesin uniquely displayed clear processive directional motion consistent with transport. These behaviours provide reference points against which the heterogeneous dynamics of HTT can be interpreted. We therefore sought to separate and quantify these motion states systematically, rather than describing HTT behaviour from population measurements alone.

### Raw-motion transformer identifies four single-molecule behavioural states with distinct frequencies across wt-HTT, Kinesin and Myosin in hESCs

To resolve heterogeneous molecular dynamics into interpretable motion states, we first manually curated visually distinct 600 ms trajectory windows and defined four recurring states: ‘bound’, ‘confined’, ‘transport-like’, and ‘exploratory’ in pluripotent undifferentiated hESCs (Figure 2A, B; Figure S3). These states capture distinct patterns of molecular motion consistent with differences in cellular interactions and physical confinement. The marked enrichment of transport-like trajectories in kinesin further supports the biological relevance of these classifications. We then quantified the curated examples using trajectory features and used PCA together with pairwise feature correlations to identify four informative non-redundant descriptors: speed, self-intersections, cumulative displacement and anomalous exponent (Figure S4).

**Figure 2.**
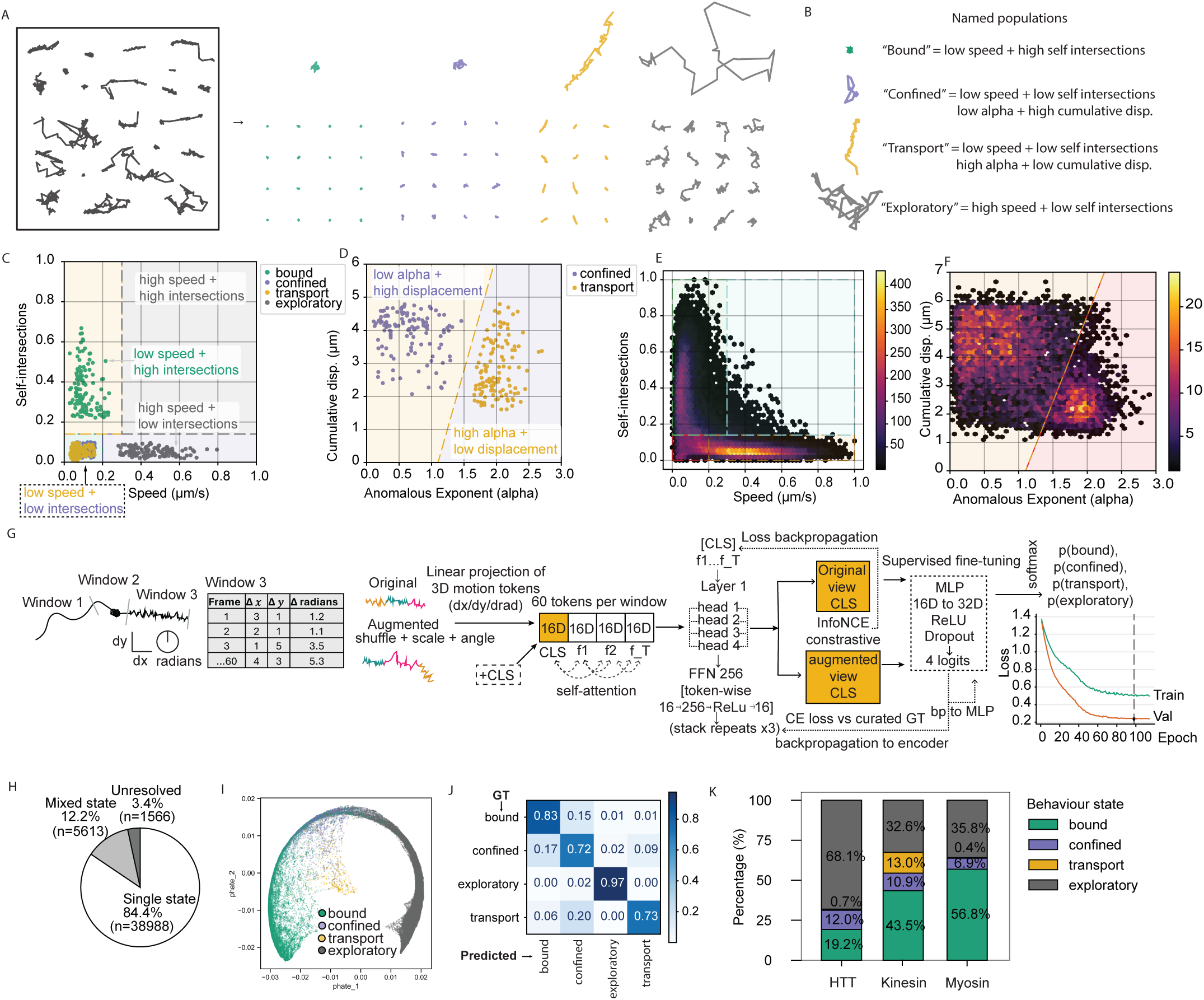
A raw-motion transformer identifies interpretable nanoscale behavioural states in single-molecule trajectories of HTT, kinesin and myosin. (A) Manually curated examples of visually distinct trajectory behaviours used to define an initial pseudo-ground-truth set. (B) Curated examples grouped into four behavioural states: bound, confined, transport-like and exploratory. (C) Mean speed versus number of self-intersections for the curated set; classification gates were defined using the separation between curated trajectory populations. (D) Cumulative displacement versus anomalous exponent for the low-speed, low-self-intersection population, distinguishing confined from transport-like trajectories. (E, F) Application of hierarchical gates to the full dataset using (E) speed and self-intersections and (F) cumulative displacement and anomalous exponent, yielding behaviour-labelled trajectory windows and revealing ambiguity near classification boundaries. (G) Schematic of the raw-motion transformer workflow. Each 600-ms trajectory window contained up to 60 ordered motion steps represented by Δx, Δy and Δθ. Each three-dimensional motion token was mapped by a learned affine projection into a 16-dimensional embedding; a learned classification (CLS) token was prepended, and the sequence was passed through a three-layer transformer encoder with four self-attention heads and a feed-forward network (16D to 256D through ReLU to 16D). Contrastive pretraining used original and augmented views of the same window as positive pairs with InfoNCE loss. Supervised fine-tuning used an MLP classification head (16D to 32D through ReLU with dropout to produce four logits) and cross-entropy loss against curated pseudo-ground-truth labels. Training and validation loss curves are shown on the right. InfoNCE loss was backpropagated through the encoder; cross-entropy loss was backpropagated through both the encoder and classifier. (H) Held-out test-set trajectory windows assigned from softmax probabilities on logits as high-confidence single-state, boundary-like two-state or unresolved low-confidence predictions. (I) PHATE projection of learned transformer embeddings, coloured by the dominant predicted behavioural state. (J) Confusion matrix comparing transformer predictions with gating-derived pseudo-ground-truth labels in a held-out test subset. (K) Transformer-defined behavioural-state distributions for HTT, kinesin and myosin in undifferentiated pluripotent hESCs.

Using these features, we constructed a hierarchical gating scheme to provide pseudo-ground-truth behavioural labels (Figure 2C-F). Bound trajectories were characterized by low speed and high self-intersections, whereas exploratory trajectories showed higher speed and fewer self-intersections. Confined and transport-like trajectories occupied relatively low-speed, low-self-intersection regions of feature space and were further separated using cumulative displacement and anomalous exponent (Figure 2C, D). Applying these gates to the full dataset generated interpretable labels, but also exposed important limitations of feature-based classification (Figure 2E, F). First, trajectories near gate boundaries were forced into discrete classes despite often exhibiting mixed or intermediate behaviour. Second, the gates depended on a small set of hand-selected summary features and dataset-specific scaling and thresholds, reducing each trajectory to averaged descriptors and discarding the temporal ordering of motion within the window. Finally, such manually defined boundaries are less readily transferable across proteins, conditions or newly acquired datasets. We therefore used the gating framework as an interpretable source of pseudo-ground-truth labels rather than as the final classifier and asked whether behavioural state could instead be inferred directly from the ordered trajectory itself.

We trained a compact raw-motion transformer using only the ordered Δx, Δy and Δθ sequence from each 600-ms trajectory window. The encoder was contrastively pretrained on augmented views of experimental trajectories and subsequently fine-tuned using the curated pseudo-ground-truth labels for bound, confined, transport-like and exploratory motion (Figure 2G; augmentation screen in Figure S5). Thus, state classification was learned from the sequence of motion itself rather than from the scalar summary features used for gating. Supervised fine-tuning used curated pseudo-ground-truth examples comprising approximately 5% of the training partition, following contrastive pretraining on trajectory sequences. A further 70% of the dataset was reserved for held-out testing and biological interpretation.

Applied to held-out hESC trajectories, the transformer returned probabilities for all four behavioural states, allowing state changes to be followed dynamically across successive trajectory windows (Movie S1: HTT, S2: kinesin, S3: myosin). Using post hoc probability criteria, 84.4% of windows received a high-confidence single state assignment, 12.2% were retained as boundary-like two-state assignments and 3.4% remained unresolved (Figure 2H). Mixed assignments preferentially occupied regions between manually defined populations in feature space, while PHATE projection of the transformer embeddings showed broad separation of the four states together with continuity between neighbouring states (Figure 2I; Figure S6). In a held-out comparison with gating-derived labels, transformer predictions agreed with the assigned labels for 83% of bound, 72% of confined, 97% of exploratory and 73% of transport trajectories (Figure 2J).

We next compared the relative abundance of these behavioural-states, hereafter behaviour state profiles, across wt-HTT, kinesin and myosin in pluripotent undifferentiated hESCs. wt-HTT was predominantly exploratory (68.1%), with smaller bound (19.2%) and confined (12.0%) populations and negligible transport-like motion (0.7%). Kinesin showed a distinct transport-like population (13.0%) together with substantial bound (43.5%), and exploratory (32.6%) and a small confined population (10.9%).

Myosin was dominated by bound trajectories (56.8%), 6.9% confined, with 35.8% exploratory and negligible 0.4% transport-like motion (Figure 2K). The enrichment of transport-like motion in kinesin was consistent with its known role in directional transport.

Together, these results establish a framework for assigning interpretable behavioural states directly from raw trajectory sequences while retaining uncertainty at class boundaries. The transformer recapitulates the distinctions defined by the curated reference behaviours without requiring hand-selected summary features for inference. Having established these behaviour-state profiles in pluripotent undifferentiated hESCs, we next applied the framework to determine how HTT, kinesin and myosin behaviours are reorganized upon neural differentiation.

### Neuronal compartmentalization differentially organizes wt-HTT, Kinesin and Myosin motion states

Neurons are highly polarized cells in which the soma, neurite and growth cone provide distinct intracellular environments. We therefore asked how wt-HTT, kinesin and myosin motion states were distributed across these compartments, focusing primarily on hESC-derived MSN-like/ventral neurons because of their disease relevance to HD (Figure 3A; Figure S7). Trajectories were segmented into 600 ms windows and classified using the raw motion transformer.

**Figure 3.**
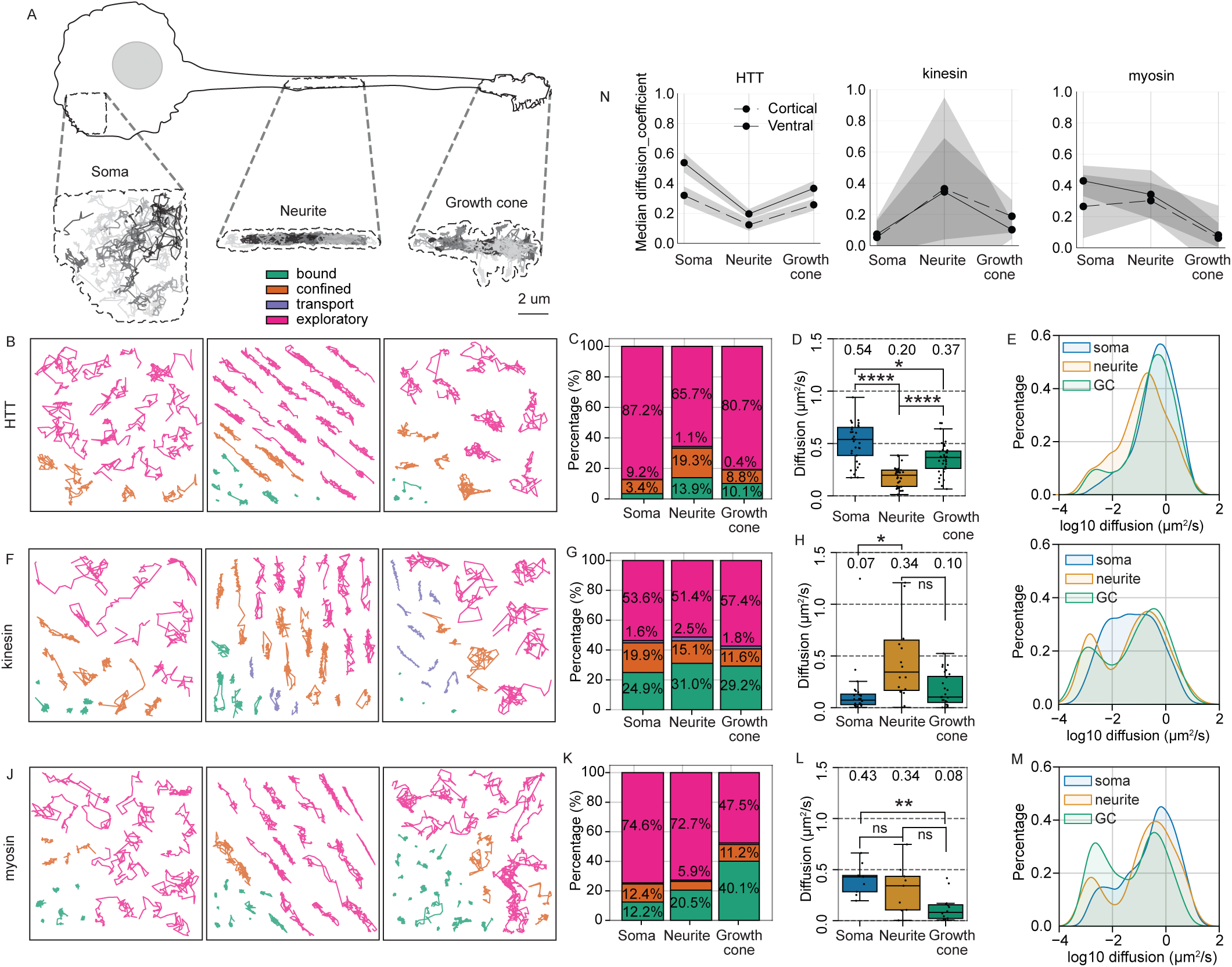
HTT, kinesin and myosin exhibit distinct compartment-specific motion profiles within ventral neurons. (A) HTT imaged in the soma, neurite and growth cone of ventral neurons. Representative trajectories are shown; contours indicate the maximum extent of trajectories within enlarged regions of interest. (B) Representative HTT trajectories in the soma, neurite and growth cone (left to right), coloured by predicted motion state. (C) Percentages of HTT trajectories assigned to each motion state in the soma, neurite and growth cone. (D) Boxplots of HTT diffusion coefficients by compartment. (E) Histograms of log10-transformed HTT diffusion coefficients by compartment. (F–I) Corresponding analyses for kinesin. (J–M) Corresponding analyses for myosin. (N) Comparison of median diffusion coefficients across neuronal compartments in ventral (solid lines) and cortical (dashed lines) neurons for HTT, kinesin and myosin. Sample sizes: HTT, 104 cells and 87,865 windowed trajectories; kinesin, 30 cells and 10,492 windowed trajectories; myosin, 30 cells and 12,463 windowed trajectories. Statistical significance was assessed using the Kruskal–Wallis test followed by Dunn’s post hoc test with Bonferroni correction. **p < 0.001, ***p < 0.0001, ****p < 0.00001; ns, not significant.

wt-HTT sampled all major behavioural states in each compartment but in different proportions (Figure 3B). Exploratory motion predominated in the soma (87.2%) and growth cone (80.7%), but decreased to 65.7% in the neurite, where bound and confined populations were more prevalent. Consistent with this compartment-dependent redistribution, median wt-HTT diffusion differed substantially across regions, measuring 0.54 µm²/s in the soma, 0.37 µm²/s in the neurite, and 0.20 µm²/s in the growth cone (Figure 3D). Diffusion coefficient distributions further revealed a shift toward slower wt-HTT motion outside the soma, with both the neurite and growth cone containing larger slow-moving subpopulations (Figure 3E). Despite the elongated geometry of neurites, most wt-HTT trajectories remained classified as exploratory rather than transport-like. Transport-like wt-HTT assignments were negligible, as in undifferentiated pluripotent hESCs, and were therefore not interpreted as evidence of processive transport.

Kinesin had a different regional profile, including a distinct transport-like population that was substantially more evident than for wt-HTT (Figure 3F). Exploratory behaviour remained relatively similar across compartments (51-57%), while substantial bound populations were also present in the soma, neurite and growth cone to similar levels (24.9-31.0%; Figure 3G). Confined behaviour was highest in the soma at 19.9%, and lower in the neurite and growth cone at 15.1% and 11.6%, respectively (Figure 3G). Transport-like trajectories were detected in all three regions and were most frequent in the neurite at 2.5 %. Kinesin mobility was also highest in the neurite, where median diffusion reached 0.34 µm²/s compared with 0.07 µm²/s in the soma and 0.10 µm²/s in the growth cone (Figure 3H). The distribution of diffusion coefficients reflected this regional shift, with the neurite showing a unimodal profile and a large mid-diffusion component, while the soma and growth cone retained a stronger low-diffusion stationary population (Figure 3I).

Myosin showed the clearest compartment-dependent shift toward stable localization, becoming markedly enriched in bound behaviour in the growth cone (Figure 3J, K). Bound behaviour increased from 12.4% in the soma and 20.5% in the neurite to 40.1% in the growth cone, accompanied by a reduction in exploratory motion to 47.5%. Median myosin diffusion correspondingly fell from 0.43 µm²/s in the soma and 0.34 µm²/s in the neurite to 0.08 µm²/s in the growth cone (Figure 3L, M). As for HTT, transport-like myosin assignments were negligible.

Despite these protein-specific distributions, wt-HTT, kinesin and myosin therefore shared the same broad repertoire of motion states across neuronal compartments but occupied them in markedly different proportions. wt-HTT was dominated by exploratory motion with greater localized confined or bound motion in the neurite and growth cone; kinesin combined exploratory and bound states with a distinct transport-associated population and maximal mobility in the neurite; and myosin display a strong enriched in immobile, bound behaviour in the growth cone. These protein-specific regional patterns were largely reproduced in cortical neurons: wt-HTT and myosin showed similar compartment-dependent changes in mobility, while kinesin again displayed its highest mobility in the neurite (Figure. 3N). Cortical and MSN-like/ventral growth cones also showed broadly similar behaviour-state profiles for all three proteins (Figure S8), indicating that the major baseline behaviours of each protein are conserved between the two neuronal identities.

### Direct effects of PolyQ expansion on the dynamics of wt-HTT protein

Having established baseline wt-HTT dynamics, we next asked whether polyQ expansion directly alters its behaviour. We generated a series of hESC lines exogenously expressing C-terminal Halo-tagged 20Q, 56Q, 77Q or 150Q HTT, alongside SNAP-tagged wt-HTT (20Q) on an HTT-/- background, with matched expressed of SNAP-versus Halo-tag in each line (Figure 4A; Figure S9). Single molecule dynamics were assessed in both undifferentiated pluripotent hESCs (Figure S10) and following their differentiation into cortical and MSN-like/ventral neurons (Figure S11). We focused initially on the growth cones of the ventral neurons because of the preferential vulnerability of the striatum in HD (Ross and Tabrizi 2011; Aylward, Nopoulos, Ross, Langbehn, Pierson, Mills, Johnson, Magnotta, Juhl, Paulsen, and Group 2011), previous evidence implicating growth-cone cytoskeletal organization in HTT function in HD (Carpentier et al. 2025), and the suitability of this dynamic, optically accessible compartment for single molecule imaging.

**Figure 4.**
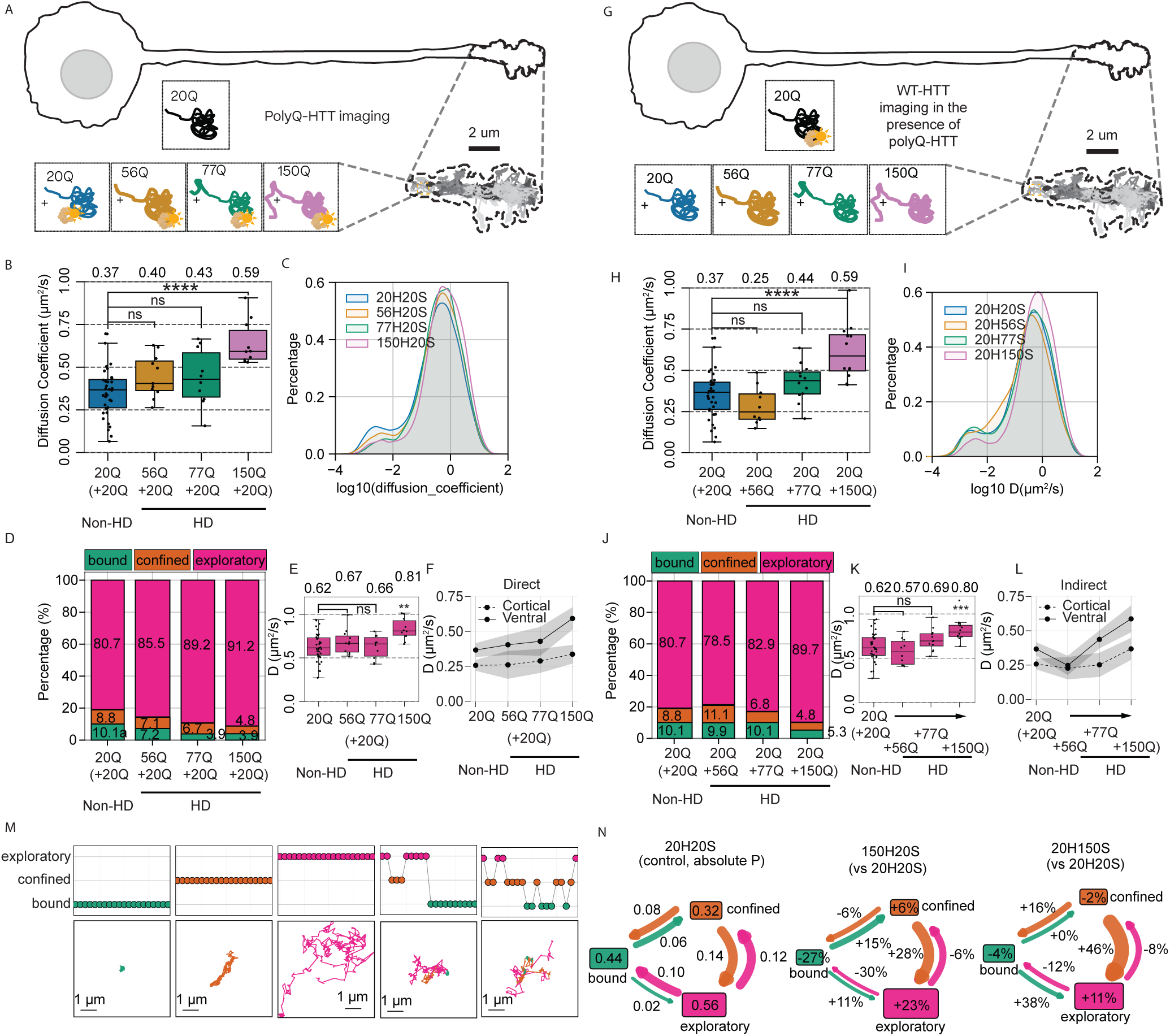
PolyQ expansion increases HTT mobility directly and through a dominant-negative effect, with ventral neurons affected at shorter PolyQ lengths than cortical neurons. (A) Schematic of the experimental design. Halo-tagged 20Q, 56Q, 77Q or 150Q HTT was imaged in ventral-neuron growth cones in the presence of SNAP-tagged 20Q HTT. (B) Boxplots of diffusion coefficients of Halo-tagged 20Q, 56Q, 77Q and 150Q HTT co-expressed with SNAP-tagged 20Q HTT in ventral-neuron growth cones. (C) Histograms of log10-transformed diffusion coefficients for each PolyQ length. (D) Behavioural-state distributions showing the percentages of trajectories assigned to each predicted motion state. (E) Diffusion coefficients of trajectories assigned to the exploratory state. (F) Comparison of median diffusion coefficients in ventral and cortical neurons. (G–L) Corresponding analyses of Halo-tagged 20Q HTT co-expressed with SNAP-tagged 20Q, 56Q, 77Q or 150Q HTT, showing the indirect effect of mutant HTT on wild-type HTT. (M) Superwindow state-transition maps and corresponding trajectories, each composed of 20 overlapping trajectory windows (nominally 6 s). (N) Fan plots of absolute state-transition probabilities for 20Q HTT (left) and percentage changes in state-transition probabilities for directly tracked 150Q HTT (middle) and 20Q HTT in the presence of 150Q HTT (right). Window counts listed for the direct series: 20Q (+20Q), 34,562; 56Q (+20Q), 10,959; 77Q (+20Q), 10,299; 150Q (+20Q), 24,306. Window counts listed for the indirect series: 20Q (+20Q), 34,562; 20Q (+56Q), 4,092; 20Q (+77Q), 7,505; 20Q (+150Q), 20,659. Combined total: 137 cells, 48,493 tracks, 146,944 windows and 5,672,760 data points. Statistical significance was assessed using the Kruskal–Wallis test followed by Dunn’s post hoc test with Bonferroni correction. **p < 0.001, ***p < 0.0001, ****p < 0.00001; ns, not significant.

HTT mobility increased progressively with PolyQ length in MSN-like/ventral growth cones, reaching a median diffusion coefficient 0.59 μm^2^/s at 150Q, compared with 0.37 μm^2^/s for 20Q, a ∼60% increase (Figure. 4B). Diffusion coefficient distributions showed a corresponding loss of the low-mobility population from 56Q onwards (Figure. 4C). Behavioural-state analysis revealed a progressive redistribution away from stable states: bound HTT fell from 10.1% at 20Q HTT to 7.2% at 56Q and 3.9% at both 77Q and 150Q, accompanied by reduced confinement and increased exploration (Figure 4D). Importantly, the median diffusion coefficient within the exploratory population remained relatively stable through 77Q, increasing only at 150Q, from 0.62 to 0.81 µm²/s relative to 20Q HTT (Figure 4E). PolyQ expansion therefore produced two distinguishable responses: 1) a progressive loss of stable bound and confined behaviour across the repeat-length series, and 2) an additional sudden increase in exploratory mobility at 150Q.

We next compared these responses with hESC-derived cortical neurons. PolyQ expansion produced the same overall direction of change in cortical growth cones but with reduced sensitivity: 56Q had little detectable effect on HTT diffusion, whereas increased mobility emerged at 77Q and was strongest at 150Q (Figure 4F; Figure S11). Thus, PolyQ-dependent disruption of HTT dynamics is not restricted to MSN-like/ventral neurons, but emerges at shorter repeat lengths in this disease-relevant neuronal lineage.

### Indirect effects of PolyQ expansion mutation in mt-HTT on the dynamics of wt-HTT proteins

We previously demonstrated a dominant-negative effect of mt-HTT on wt-HTT activity in hESC micropattern and neuruloid models (Laundos et al. 2023). Here, we asked whether this effect could also be detected in the dynamics of individual wt-HTT single molecules. To address this, we reversed the tags: wild-type 20Q HTT was Halo-tagged and tracked in the presence of SNAP-tagged 20Q, 56Q, 77Q or 150Q HTT (Figure 4G). In MSN-like/ventral growth cones, wt-HTT mobility was unchanged in the presence of co-expressed 56Q mt-HTT, but was increased with 77Q and further increased with 150Q, reaching 0.59 µm²/s; a 59.5% increase from 0.37 µm²/s in the wt-HTT 20Q/20Q condition (Figure 4H). Diffusion coefficient distributions showed a shift from lower toward higher mobility (Figure 4I), analogous to the direct effect of the mutation (Figure 4C). Thus, mt-HTT strongly alters the dynamics of wt-HTT molecules sharing the same cellular environment.

Behavioural state analysis ratified this dominant effect. Co-expression with 56Q or 77Q mt-HTT produced little change in the overall behaviour-state profile of wt-HTT, whereas 150Q caused a clear loss of bound behaviour from 10.1 to 5.3% (almost half) and confined behaviour from 8.8 to 4.3%, accompanied by increased exploration (Figure 4J). Mobility within the exploratory population also increased in the presence of 150Q mt-HTT, reaching 0.80 μm^2^/s compared with 0.62 um^2^/s in the wt-HTT 20Q/20Q control (Figure 4K). The dominant effect therefore comprises both redistribution between behavioural states and increased mobility within the exploratory state, but these readouts emerge with different PolyQ-length dependence.

This dominant effect also differed between neuronal identities. In cortical neurons, increased mobility of wt-HTT was detected only in the presence of 150Q mt-HTT, whereas in ventral/MSN-like neurons it was already apparent with 77Q (Figure 4L). Across soma, neurite and growth cone, mt-HTT altered wt-HTT dynamics in both neuronal types, but the effects were most pronounced in the growth cone (Figure S10). These findings further indicate greater sensitivity of ventral/MSN-like neurons to the molecular consequences of PolyQ expansion. A similar combination of direct and indirect dominant effects was observed in undifferentiated hESCs and cortical neurons but was less pronounced and emerged predominantly at larger PolyQ tracts (Figure S10, S11).

### PolyQ expansion disrupts wt-HTT behavioural-state switching

Having shown that PolyQ expansion changes the proportion of HTT molecules occupying each behavioural state, we next asked whether it also alters how individual molecules switch between states over longer timescales. We therefore followed consecutive transformer-defined behaviour states within longer trajectories and quantified the probability that wt-HTT either remained in its current state or transitioned to another (Figure 4M; additional trajectories in Figure S12; example tracks in Movie S1).

In 20Q wt-HTT, exploratory behaviour was the most persistent of the three major states with P(remain) 0.56, while bound (0.44) and confined (0.32) molecules showed greater state switching probability. Transitions occurred most frequently between confined and exploratory (P(switch) 0.12 or 0.14), whereas direct transitions from bound to exploratory motion were comparatively rare (P(switch) 0.02; Figure 4N). Thus, wt-HTT dynamically moves between localized and exploratory modes while retaining a measurable propensity to maintain stable engagement.

In mt-HTT protein carrying 150Q expansion, this balance shifted markedly toward exploratory behaviour. Relative to wt-HTT, mt-HTT was 27% less likely to remain bound and 23% more likely to remain exploratory. Transitions from confined (+28%) or bound (+11%) states toward exploration increased, whereas the probability of transitioning from exploratory motion back to the bound state decreased by 30% (Figure 4N; middle fan diagram). PolyQ expansion therefore affects not only state occupancy but also behavioural plasticity, destabilizing persistent engagement and biasing wt-HTT toward exploratory motion.

20Q wt-HTT co-expressed with 150Q mt-HTT showed a related but distinct switching phenotype. Its probability of remaining bound was only modestly reduced, whereas transitions from confined or bound states into exploratory motion increased substantially, while return from exploration to binding was reduced (Figure 4N). Thus, pronounced loss of bound-state maintenance was primarily a direct property of mt-HTT, whereas presence of HD mutation also biased both mt-and co-expressed wt-HTT molecules toward exploratory transitions. Transition probabilities across the complete PolyQ series are shown in Figure S13.

### Both mt-HTT and HTT-/- disrupt kinesin and myosin dynamics in neuronal growth cones

The dominant effect of mt-HTT on co-expressed wt-HTT suggested that PolyQ expansion alters the intracellular environment in which wt-HTT operates. To test whether this extends to other proteins, we measured kinesin and myosin dynamics in MSN-like/ventral growth cones expressing wt-HTT, endogenous mt-HTT carrying 72Q, or HTT-/- (Figure 5A). Kinesin mobility increased markedly in both mt- and HTT-/- cells, from a median diffusion coefficient of 0.14 um^2^/s in wild-type to 0.36 um^2^/s with mt-HTT (more than 2-fold increase) and 0.30 um^2^/s in HTT-/- cells (2.1 fold increase; Figure 5B). Diffusion-coefficient distributions showed increased high-mobility kinesin and reduced low- and intermediate-mobility populations in both genetic perturbations (Figure 5C).

**Figure 5.**
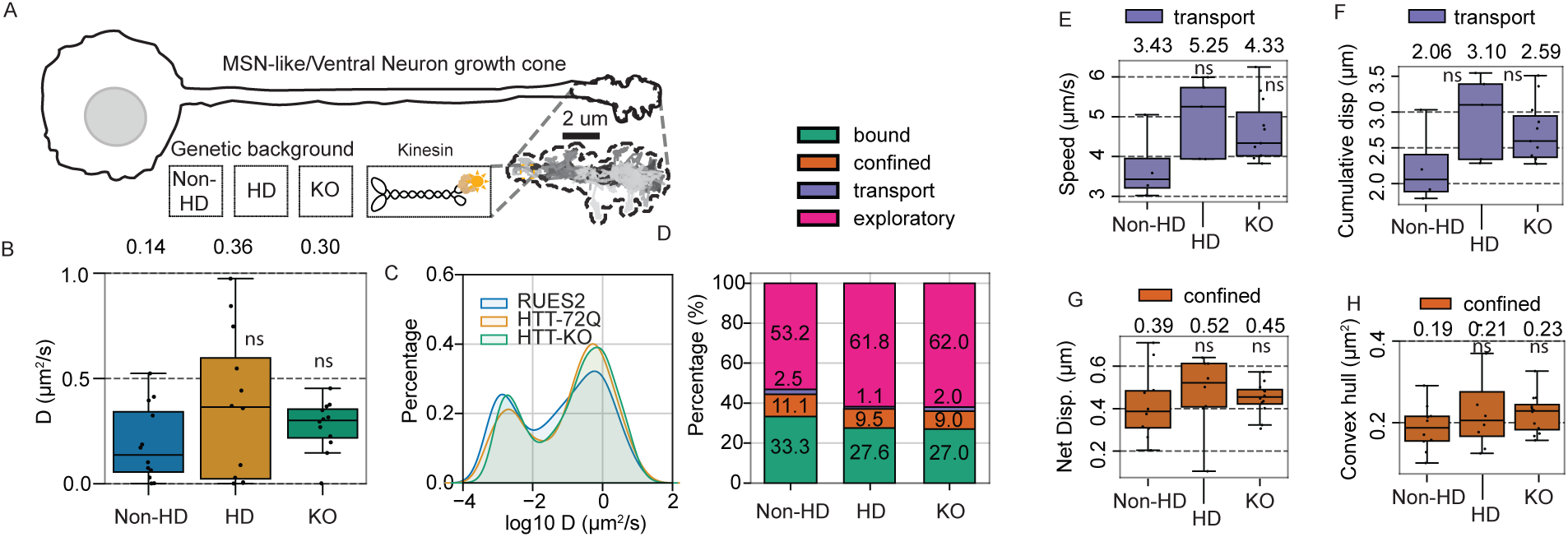
HTT mutation or loss shifts kinesin from bound, confined and transport-like states toward exploratory motion. (A) Schematic of Halo-tagged kinesin imaging in the growth cones of ventral neurons with wild-type HTT (HTT-WT), polyQ-expanded HTT (HTT-72Q) or HTT knockout (HTT-KO) backgrounds. (B) Boxplots of kinesin diffusion coefficients in each HTT background. (C) Histograms of log10-transformed kinesin diffusion coefficients in each HTT background. (D) Behavioural-state distributions showing the percentages of trajectories assigned to each predicted motion state. (E) Speeds of trajectories assigned to the transport-like state. (F) Cumulative displacements of trajectories assigned to the transport-like state. (G) Net displacements of trajectories assigned to the confined state. (H) Convex hull areas of trajectories assigned to the confined state. Sample sizes: HTT-WT, 12 cells and 8,638 windowed trajectories; HTT-72Q, 11 cells and 10,054 windowed trajectories; HTT-KO, 12 cells and 7,644 windowed trajectories. Statistical significance was assessed using the Kruskal–Wallis test followed by Dunn’s post hoc test with Bonferroni correction.

Behavioural state analysis showed that this increased mobility reflected a redistribution of kinesin away from bound and confined states toward exploratory motion (Figure 5D). The transport associated population also decreased from 2.5% in wild-type to 1.1% with mt-HTT and, more modestly to 2.0%, in HTT-/- cells. However, among kinesin trajectories that remained transport-like, movement was faster, increasing from 3.43 μm/s in wt-to 5.25 μm/s with mt-HTT and 4.33 μm/s in HTT-/- cells (Figure 5E). A similar combination of reduced transport-state occupancy and increased speed within the remaining transport population was observed in undifferentiated pluripotent hESCs (Figure. S14). Thus, increased overall kinesin mobility is associated primarily with loss of stable localization/binding and increased exploration, while the smaller population that remains transport-like moves faster, despite being less frequent.

Kinesin dynamics were also altered within the confined state. Such confinement may reflect transient interactions with intracellular structures or physical restriction within locally organized domains. Confined trajectories showed increased convex-hull area and net displacement in both mt- and HTT-/- growth cones, indicating that kinesin explored larger, less tightly restricted domains, despite remaining within the same behavioural class (Figure 5G, H). The same effect was observed in undifferentiated hESCs (Figure. S14), providing further evidence that HD mutation or loss of HTT alters the local environment governing kinesin localization and confinement.

Myosin responded more selectively. mt-HTT produced relatively little change in myosin dynamics in ventral/MSN-like growth cones, whereas HTT-/- produced a stronger effect; both perturbations altered myosin dynamics strongly in pluripotent undifferentiated hESCs (Figure S15). Thus, disruption of HTT function alters the dynamics of other cytoskeletal-associated proteins, but with protein-and cellular-context dependent effects that are substantially stronger for kinesin in neuronal growth cones.

Together, these results show that both mutant HTT and loss of HTT remodel kinesin behaviour, reducing stable localization and transport-associated occupancy while increasing exploratory motion and the spatial range of confined trajectories. The related effects on myosin further support alteration of the intracellular environment in which these proteins operate.

## DISCUSSION

### HTT dynamics at single-molecule resolution

The motion of a protein within a cell reflects its interactions with, and physical constraints imposed by, the surrounding intracellular environment. Single-molecule tracking therefore provides an indirect readout of molecular engagement(Chen et al. 2014; Liu et al. 2015; Piccolo et al. 2019; Shannon et al. 2020). Stable localization may reflect sustained interactions with immobile elements, confinement may indicate movement within spatially restricted domains, and exploratory motion may represent movement between sites of engagement. These behaviours are consistent with molecular engagement or physical restriction, but motion alone cannot establish a biochemical interaction or identify the partner involved. These behaviours should not be interpreted as fixed molecular identities, but as states that individual molecules can enter and leave over time. Tracking their occupancy over time and their interconversion can therefore reveal how readily a protein such as HTT engages with, disengages from, and returns to localized intracellular environments, even when the molecular partner responsible for each state remains unknown.

Machine-learning approaches increasingly extract temporal information directly from single-particle trajectories. DeepSPT identifies canonical diffusion modes and their transitions (Kæstel-Hansen et al. 2025), whereas SPTnet jointly reconstructs trajectories and estimates physical motion parameters (Bi et al. 2025). Our approach addresses a complementary question: rather than assigning predefined diffusion modes, it learns experimentally defined, protein-relevant behaviours from raw trajectory sequences. As in previous work on cell level behaviour tracing (Shannon et al. 2024), we focused on making the analysis interpretable by relating model predictions to experimentally recognizable behaviours, rather than treating the classes identified by the model as biological findings in themselves. This allowed us to analyse large numbers of trajectories while preserving their connection to observable molecular behaviour. The resulting framework allowed bound, confined, exploratory and transport-like motion to be compared consistently across proteins and genotypes, while retaining intermediate assignments between states.

Following these behaviours over longer trajectories extends the analysis from state occupancy to behavioural plasticity: the ability of molecules to enter, maintain and leave particular modes of motion. The framework is also extensible: additional experimentally validated behaviours can be introduced as additional reference states, providing a route from molecular trajectories toward functional interpretation of disease associated changes.

### HTT occupies a dynamic regime distinct from those of kinesin and myosin

This study resolves the behaviour of individual full-length HTT molecules in living human pluripotent cells and hESC-derived cortical and MSN-like/ventral neurons at high temporal resolution. Using this method, we found that HTT was exclusively cytosolic in pluripotent cells, neuronal soma, neurites, and growth cones, with no detectable nuclear localization events. These observations establish a live-cell baseline in which full-length HTT is predominantly a mobile cytoplasmic protein whose behaviour reflects repeated transitions between exploratory, confined and stably localized states. We found that HTT dynamics were not influenced by expression level, in agreement with the wide range of expression levels that has been reported for HTT in the human population (Fodale et al. 2022).

Comparison with kinesin and myosin places this HTT behaviour in a functional context, as both are well-characterized motor proteins that interact differentially with the cytoskeleton and can therefore act as functional reference points for interpreting HTT motion. HTT has long been implicated in cytoskeleton-dependent processes, including vesicular transport along microtubules and more recently the organization of both microtubule and actin-associated structures (Gauthier et al. 2004; Caviston et al. 2007; Capizzi et al. 2022; Carpentier et al. 2025). Across undifferentiated hESCs and neurons, HTT was substantially more mobile than either protein and was predominantly exploratory, whereas myosin showed the strongest enrichment of bound behaviour and kinesin displayed the clearest transport-associated population. Thus, although all three proteins sampled the same broad repertoire of motion states, they occupied those states in different proportions. HTT did not display the strongly localized signature characteristic of myosin (Quintanilla et al. 2023), nor a substantial processive transport population comparable with kinesin (Verhey and Ohi 2023). In particular, transport-like HTT assignments remained negligible across the cellular contexts examined, arguing against processive movement being a substantial behaviour of full-length HTT under these conditions. Instead, HTT predominantly sampled exploratory motion, punctuated by periods of confinement and stable localization.

### PolyQ expansion directly and indirectly destabilizes HTT dynamics

PolyQ expansion increased the mobility of full-length HTT, which initially appears to contrast with the literature emphasizing PolyQ-dependent HTT misfolding, aggregation and inclusion-body formation in HD (Arrasate and Finkbeiner 2012). However, these observations are not mutually exclusive. Our hESC-derived neurons are developmentally young, and imaging at high temporal resolution enables resolution of the behaviour of the highly mobile HTT population, revealing rapid changes in molecular engagement that are inaccessible to aggregate-based or endpoint measurements. Such changes may therefore represent earlier molecular dysfunction, before visible inclusion formation becomes a dominant phenotype.

Fast imaging at single-molecule resolution allowed us to distinguish two distinct PolyQ expansion-dependent changes in HTT behavior. First, increasing repeat length progressively shifted mutant HTT away from bound and confined states toward exploratory motion, consistent with reduced stable engagement. Second, mobility within the exploratory population remained relatively stable through 77Q but increased at 150Q. Thus, progressive disruption of HTT engagement coexists with an additional long-repeat-length-specific phenotype, compatible with both progressive dysregulation of HTT behaviour with PolyQ expansion as well as a threshold phenotype appearing only at the higher repeat length, both shown for the first time at single molecule resolution. Either effect has been suggested as an alternative consequence of somatic expansion in HD patients, while our data suggest that the two may be reconcilable (Mätlik et al. 2024; Handsaker et al. 2025).

The PolyQ-dependent phenotype emerged at shorter repeat lengths in MSN-like/ventral neurons than in cortical neurons or hESCs and was strongest in the growth cone. Because wild-type cortical and MSN-like/ventral growth cones showed broadly similar baseline HTT dynamics, this earlier response might reflect a greater sensitivity to PolyQ expansion rather than a pre-existing difference in HTT mobility, consistent with the preferential vulnerability of striatal MSNs in HD (Ehrlich 2012, Ryoma and Goto 2017).

The tag-swap experiments showed that PolyQ expansion also alters the dynamics of co-expressed wild-type HTT. In the presence of mutant HTT, wild-type HTT became more mobile and, at longer PolyQ lengths, shifted away from bound and confined states, indicating a loss of localization and binding, broadly mirroring the direct mutant phenotype and supporting a dominant effect of HD mutation (Laundos et al. 2023). However, the two effects were not identical. Measuring state switching/plasticity over longer timescales revealed that while mutant 150Q HTT itself showed a pronounced loss of bound-state stability when already in a bound state, whereas this was only modestly reduced in wild-type HTT exposed to the same 150Q environment. Instead, wild-type HTT became more likely to transition toward exploration (from bound or confined) and less likely to return to binding. Thus, direct and indirect effects occur alongside one another, but stable binding is more severely disrupted in mutant HTT itself.

### HD mutation or loss of HTT expression disrupts the intracellular environment

The dominant effect on wild-type HTT could arise through HTT-HTT association or alteration/deterioration of their shared intracellular environment. Co-immunoprecipitation experiments did not detect stable associations between differentially tagged WT-WT, mutant-mutant or WT-mutant HTT under the conditions tested (Figure S16). These findings argue against stable HTT–HTT association as the principal explanation for the observed dominant dynamic phenotype, although transient interactions cannot be excluded. This then led us to ask whether HD mutation instead alters the behaviour of other proteins within the same environment. Kinesin provided the clearest evidence that this shared environment is disrupted by both HD mutation and HTT loss. In both undifferentiated hESCs and MSN-like/ventral neurons, these perturbations increased kinesin mobility and shifted molecules away from bound and confined states toward exploration. The transport-like population decreased, while the remaining transport-like molecules moved faster, suggesting that fewer kinesin trajectories exhibited sustained directional motion, although those that did may have experienced reduced mechanical constraint or altered regulation. Furthermore, confined kinesin explored larger areas and underwent greater net displacement, consistent with looser spatial restriction and a less tightly organized cytoskeletal meshwork. This may reflect changes in the physical constraints imposed by intracellular structures, conceptually similar to the compartmentalized diffusion described by the picket-fence model (Kusumi et al. 2005). Related changes in myosin suggest that this effect is not restricted to kinesin. Together with previous work linking HTT to cytoskeletal organization and motor regulation/trafficking (Gauthier et al. 2004, Caviston et al 2007, Colin et al. 2008, Juliane and Holzbaur 2009, Capizzi et al 2022, Carpentier et al. 2025,) and with current investigation in our laboratory on altered microtubule dynamic instability in HD and HTT-KO cells (Kim et al. in preparation), these findings suggest that HTT helps organize the local environment in which motor proteins, and HTT itself, are positioned and engage with the cytoskeleton. The similarity between HD and HTT-KO therefore supports a loss-of-normal-function component in which this organization becomes disrupted.

### Limitations of the study

Overall, this study defines the dynamic behaviour of full-length HTT at single-molecule resolution in living human cells and shows that PolyQ expansion alters the dynamic interactions of both HTT itself and the intracellular environment in which it operates. Together with complementary structural observations from our group (Kim et al., in preparation), these findings support a role for HTT in cytoskeletal organization. The molecular mechanisms through which HTT contributes to this organization, how PolyQ-expanded HTT disrupts it, and how these nanoscale dynamic changes ultimately contribute to neuronal dysfunction and the neurodegeneration observed in HD patients remain to be established. Resolving these links will be important for connecting the molecular behaviour of HTT to the cellular pathogenesis of Huntington’s disease.

## RESOURCE AVAILABILITY

### Lead contact

Further information and requests for resources and reagents should be directed to and will be fulfilled by the lead contact, Ali Brivanlou.

### Materials availability

Cell lines and plasmids generated in this study are available from the lead contact upon reasonable request and completion of an appropriate material transfer agreement, where required.

### Data and code availability

Single-particle tracking data generated in this study will be deposited and will be publicly available as of the date of publication. SPTnano analysis workflow, trained transformer model, preprocessing code and associated analysis tools are publicly available through GitHub (https://github.com/Michael-shannon/SPTnano-core).

### Declaration of generative AI and AI-assisted technologies in the manuscript preparation process

During the preparation of this work, the author(s) used Claude for coding assistance. The author(s) reviewed and edited the output as needed and take full responsibility for the content of the published article.

## ACKNOWLEDGMENTS

We thank some people for technical assistance/discussions/reagents, and members of the Brivanlou laboratory for helpful discussions. We acknowledge FACS facility, Bio-Imaging Resource Center (BIRC) and High Performance Computing facility at Rockefeller University for assistance with TIRF imaging, FACS and Compute. This work was supported by Cure Hungtington’s Disease Initiative (CHDI).

## AUTHOR CONTRIBUTIONS

Conceptualization, MS, FP; Methodology, MS, FP; Software, MS; Validation, MS; Formal Analysis, MS; Investigation, MS, SL, TN, TLL, CK; Resources, AHB, FP, SL, TN, TLL; Data Curation, MS; Original Draft, MS; Review & Editing, MS, SL, TN, TLL, CK, FP, AHB; Visualization, MS; Supervision, AHB, FP; Project Administration, AHB, FP; Funding Acquisition, AHB, FP.

## STAR Methods

### Key Resources Table

| **Reagent/Resource** | **SOURCE** | **IDENTIFIER** |
|---|---|---|
| **Experimental models: Cell lines** |  |  |
| RUES2 human embryonic stem cells | NIH Human Embryonic Stem Cell Registry | RUES2-NIHhESC-09-0013 |
| HTT-knockout RUES2 hESCs | Ruzo et al., 2018 / Brivanlou laboratory | HTT-KO RUES2 |
| Endogenous HTT-HaloTag RUES2 reporter line | Laundos et al., 2023 / Brivanlou laboratory | HTT-HaloTag RUES2 |
| 20QHTT-Halo/20QHTT-SNAP RUES2 hESCs | This study | 20H20S |
| 56QHTT-Halo/20QHTT-SNAP RUES2 hESCs | This study | 56H20S |
| 77QHTT-Halo/20QHTT-SNAP RUES2 hESCs | This study | 77H20S |
| 150QHTT-Halo/20QHTT-SNAP RUES2 hESCs | This study | 150H20S |
| 20QHTT-Halo/56QHTT-SNAP RUES2 hESCs | This study | 20H56S |
| 20QHTT-Halo/77QHTT-SNAP RUES2 hESCs | This study | 20H77S |
| 20QHTT-Halo/150QHTT-SNAP RUES2 hESCs | This study | 20H150S |
| KIF5B-HaloTag RUES2 hESCs, WT HTT background | This study | KIF5B-Halo WT |
| KIF5B-HaloTag RUES2 hESCs, HTT-72CAG background | This study | KIF5B-Halo HTT-72CAG |
| KIF5B-HaloTag RUES2 hESCs, HTT-KO background | This study | KIF5B-Halo HTT-KO |
| MYH9-HaloTag RUES2 hESCs, WT HTT background | This study | MYH9-Halo WT |
| MYH9-HaloTag RUES2 hESCs, HTT-72CAG background | This study | MYH9-Halo HTT-72CAG |
| MYH9-HaloTag RUES2 hESCs, HTT-KO background | This study | MYH9-Halo HTT-KO |
| **Recombinant DNA** |  |  |
| PiggyBac full-length HTT-20-SNAP | This study | N/A |
| PiggyBac full-length HTT-20-HaloTag | Laundos et al. 2023 | N/A |
| PiggyBac full-length HTT-56-SNAP | This study | N/A |
| PiggyBac full-length HTT-56-HaloTag | Laundos et al. 2023 | N/A |
| PiggyBac full-length HTT-77-SNAP | This study | N/A |
| PiggyBac full-length HTT-77-HaloTag | This study | N/A |
| PiggyBac full-length HTT-150- SNAP | This study | N/A |
| PiggyBac full-length HTT-150-HaloTag | This study | N/A |
| PiggyBac KIF5B C-term HaloTag | This study; from KIF5B (Addgene 15219) | N/A |
| PiggyBac MYH9 C-term HaloTag | This study; from MYH9 (Addgene 38297) | N/A |
| PiggyBac transposase | Lacoste et al., 2009 | N/A |
| **Chemicals, peptides, and recombinant proteins** |  |  |
| Geltrex LDEV-Free, hESC-Qualified, Reduced Growth Factor Basement Membrane Matrix | Thermo Fisher | A1413302 |
| Accutase Cell Detachment Solution | StemCell Technologies | 07920 |
| Janelia Fluor HaloTag Ligand, JFX 554 | Promega | HT1030 |
| Nucleofector Kit L | Lonza | VCA-1005 |
| bFGF | PeproTech | PHG0263 |
| GDNF | R&D Systems | 212-GD-050/CF |
| BDNF | R&D Systems | 248-BDB-050/CF |
| FGF-8b | R&D Systems | 423-F8-025 |
| IGF1 | R&D Systems | 291-G1-200 |
| DAPT | Millipore Sigma | D5942-5MG |
| Dibutyryl-cAMP | R&D Systems | 1141 |
| Laminin-521 | Biolamina | LN521-05 |
| N2 supplement | Thermo Fisher | 17502048 |
| B27 supplement with vitamin A | Life Technologies | 17504-044 |
| B27 supplement without vitamin A | Gibco | 12587-010 |
| **Software and algorithms** |  |  |
| Slimfast | Wong et al., 2025 | DOI: 10.5281/zenodo.14901152 |
| SPTnano | This study | https://github.com/Michael-shannon/SPTnano |
| Python | Python Software Foundation | v3.10–3.12; RRID:SCR_008394 |
| NumPy | Harris et al., 2020 | ≥1.24; RRID:SCR_008633 |
| pandas | The pandas development team | ≥2.0; RRID:SCR_018214 |
| Polars | Polars developers | https://github.com/pola-rs/polars |
| SciPy | Virtanen et al., 2020 | ≥1.10; RRID:SCR_008058 |
| joblib | Joblib developers | ≥1.3; https://joblib.readthedocs.io |
| PyTorch | Paszke et al., 2019 | ≥2.0; RRID:SCR_018536 |
| scikit-learn | Pedregosa et al., 2011 | ≥1.3; RRID:SCR_002577 |
| UMAP (umap-learn) | McInnes et al., 2018 | ≥0.5; https://github.com/lmcinnes/umap |
| HDBSCAN | McInnes et al., 2017 | ≥0.8.29; https://github.com/scikit-learn-contrib/hdbscan |
| matplotlib | Hunter, 2007 | ≥3.7; RRID:SCR_008624 |
| seaborn | Waskom, 2021 | ≥0.13; RRID:SCR_018132 |
| napari | Sofroniew et al. | RRID:SCR_022765; https://napari.org |
| scikit-image | van der Walt et al., 2014 | ≥0.21; RRID:SCR_021142 |
| Jupyter / JupyterLab | Kluyver et al., 2016 | RRID:SCR_018315 |
| SciPy Dunnett test (scipy.stats.dunnett) | Virtanen et al., 2020 | SciPy ≥1.11, RRID:SCR_008058 |

### EXPERIMENTAL MODEL AND STUDY PARTICIPANT DETAILS

#### Human embryonic stem cell culture

Human embryonic stem cell lines used in this study were derived from the RUES2 cell line, which is registered in the NIH Human Embryonic Stem Cell Registry (RUES2-NIHhESC-09-0013). hESCs were maintained under feeder-free conditions in mouse embryonic fibroblast-conditioned HUES medium supplemented with 20 ng/mL bFGF, with medium replaced daily. Cells were passaged at 70% to 80% confluency every 3 to 4 days onto Geltrex-coated dishes (Life Technologies) using Gentle Cell Dissociation Reagent (Stemcell Technologies). Cultures were tested for *Mycoplasma* spp. at 2-month intervals.

#### Cortical and ventral/MSN-like neuronal differentiation

RUES2 human embryonic stem cells were differentiated into cortical- and ventral-patterned neuronal cultures using a staged neural induction, neural progenitor and neuronal maturation protocol adapted from established dual-SMAD/WNT-inhibition approaches (Qi et al., 2017). hESCs were seeded 2 days before neural induction onto Geltrex-coated dishes at 5 × 10⁴ cells/cm² and allowed to reach confluence before differentiation. From day 0 to day 2, neural induction was performed in Essential 6-based medium containing 100 nM LDN193189, 10 µM SB431542 and 2 µM XAV939 (LSBX). XAV939 was withdrawn from day 3, and cultures were maintained with 100 nM LDN193189 and 10 µM SB431542 through day 7. From day 8 to day 11, cultures were transitioned to N2/B27-based neural progenitor medium comprising a 1:1 mixture of DMEM/F12 and Neurobasal medium supplemented with 0.5× N2 (Gibco/Thermo Fisher Scientific, 17502048), 0.5× B27 without vitamin A (Gibco/Thermo Fisher Scientific, 12587010), β-mercaptoethanol, insulin, GlutaMAX and non-essential amino acids. From day 12, B27 without vitamin A was replaced by 0.5× B27 containing vitamin A (Gibco/Thermo Fisher Scientific, 17504044), and cultures were supplemented with 0.2 mM ascorbic acid, 0.5 mM dibutyryl-cAMP (dbcAMP; Tocris Bioscience/Bio-Techne, 1141), 20 ng/mL BDNF (R&D Systems, 248-BDB-050/CF) and 100 ng/mL FGF8b (R&D Systems, 423-F8-025). Cells were replated at day 20 on 10 ug/ml laminin-521 (BioLamina, LN521-05) coated dishes in maturation medium containing ascorbic acid, dbcAMP, BDNF and FGF8b, with 10 µM Y-27632 included during replating. FGF8b was subsequently withdrawn, and cultures were maintained with 0.2 mM ascorbic acid, 0.5 mM dbcAMP and 20 ng/mL BDNF, with half-medium changes performed every other day. For cortical differentiation, 10 µM DAPT (MilliporeSigma, D5942) was added from day 20 to day 30 to promote neuronal differentiation. After day 30, cortical maturation medium was additionally supplemented with 20 ng/mL GDNF (R&D Systems, 212-GD-050/CF). Cortical cultures were cryopreserved at day 20. For experiments, day-20 cortical stocks were thawed, matured further with half-medium changes every other day, and used for imaging at approximately day 35. For ventral/MSN-like differentiation, Smoothened agonist (SAG; 0.1 µM) was added from day 5 through day 25 to promote ventral patterning. DAPT was not used in ventral cultures. From day 31 onwards, ventral maturation medium was supplemented with 20 ng/mL GDNF in addition to 0.2 mM ascorbic acid, 0.5 mM dbcAMP and 20 ng/mL BDNF. Ventral neuronal cultures were cryopreserved at day 35. For experiments, day-35 ventral stocks were thawed, matured further with half-medium changes every other day, and used for imaging at approximately day 45. Neuronal differentiation and regional identity were verified by marker expression using 1:5000 MAP2 (Abcam, ab5392), 1:250 DARPP32 (Abcam, ab40801), 1:1000 CTIP2 (Abcam, ab18465), confirming the generation of cortical- and ventral/MSN-like neuronal populations (Figure S7].

#### Generation of full-length HTT HaloTag- and SNAP-tagged hESC lines

HTT-knockout RUES2 hESCs were nucleofected using an Amaxa Nucleofector 2 with Nucleofector Solution L (Lonza) and program B-016. For each nucleofection, cells were transfected with 1 μg PiggyBac full-length HTT-HaloTag plasmid and 1 μg PiggyBac transposase (Lacoste et al. 2009). Following nucleofection, cells were cultured in conditioned medium supplemented with 20 ng/mL bFGF and 10 μM Y-27632 for 24 h and subsequently selected with 1 μg/mL puromycin or 10 µg/mL blasticidin. Stable expression was confirmed before fluorescence-activated cell sorting. For double tagging, sequential nucleofection selection cycles were used. Full-length HTT constructs contained CAG repeat lengths encoding polyglutamine tracts of 20Q, 56Q, 77Q, or 150Q. HaloTag- and SNAP-tagged versions of the constructs were co-expressed to generate the following cell lines, where H denotes the HaloTag-labelled HTT allele and S denotes the SNAP-tagged HTT allele: 20H20S, 56H20S, 77H20S, 150H20S, 20H56S, 20H77S, and 20H150S. HaloTag and SNAP-tag expression levels were assessed, and populations with comparable expression of the two tagged constructs were isolated by fluorescence-activated cell sorting as described below.

#### Generation of endogenous HaloTag reporter cell line

An endogenous HTT-HaloTag reporter RUES2 line carrying a monoallelic C-terminal HaloTag fusion at the endogenous HTT locus was generated by CRISPR-mediated genome editing and was previously used as an endogenous-expression reference (Laundos et al. 2023). The reporter was validated for correct HTT-HaloTag expression by western blot and comparison to exogeneously expressed HTT-HaloTag with single particle tracking (Figure S1).

#### Generation of KIF5B-HaloTag and MYH9-HaloTag lines on WT, HTT-72CAG and HTT-KO backgrounds

The HTT-72CAG line was derived from a previously described isogenic RUES2 HD PolyQ series, in which one endogenous HTT allele carries an expanded CAG tract; the HTT-KO line was also derived as previously described (Ruzo et al. 2018). A truncated version of KIF5B, consisting of the first 560 amino acids (derived from Addgene plasmid #15219) was fused to HaloTag at the C-terminus in a PiggyBac backbone. Full-length MYH9 (derived from Addgene plasmid #38297) was also HaloTagged at the C-terminus in a PiggyBac backbone. 1 μg of each plasmid was nucleofected alongside 1 μg of PiggyBac Transposase into WT, HTT-72CAG and HTT-KO backgrounds, then selected and maintained as described above. Cell population expressing low levels of HaloTag was isolated by FACS. HaloTag expression patterns was cross-checked with endogenous KIF5B and MYH9 expression by immunofluorescence (Figure S2).

#### Fluorescence-activated cell sorting

Cells below confluency were incubated for 1 h at 37°C in conditioned medium supplemented with bFGF and 1 μM cell-permeable Janelia Fluor X 554 HaloTag ligand (Promega, GA1110). Cells were washed three times with PBS+/+ and incubated in fresh medium for 3 h to allow removal of unbound ligand. Monolayers were subsequently washed with PBS−/−, dissociated using Accutase, and resuspended in PBS−/− containing 10 mM HEPES, 5 mM EDTA, and 0.5% BSA. Doublets were excluded using FSC-W and SSC-W gating, and DAPI was included at 20 ng/mL for exclusion of non-viable cells. Cells were sorted using a BD FACSAria flow cytometer with a 100-μm nozzle at 20 psi. The parental untagged cell line was treated with HaloTag ligand in parallel and used as a negative control. Following sorting, cells were allowed to recover in medium containing Normocin. Sorted populations were subsequently evaluated by flow cytometry, western blotting, and single-particle measurements.

#### Preparation of cells for imaging

For hESC imaging, cells were transferred to four-well glass-bottom Ibidi dishes coated with laminin-521 (10 μg/mL in PBS+/+) for 2 h at 37°C. Cells were maintained for 12 h in conditioned HUES medium supplemented with 20 ng/ml bFGF and 10 μM Y-27632. Medium was then replaced, and cells were cultured for a further 24 h before labelling. JFX-554 HaloTag ligand was added at 0.5 nM in phenol-red-free conditioned medium for 15 min, followed by one wash in the same medium before imaging. For cortical and ventral/MSN-like neurons, day-20 or day-35 cryopreserved cultures, respectively, were thawed onto laminin-521-coated four-well glass-bottom dishes and maintained in the corresponding neuronal differentiation medium with medium changes every 48 h. Before imaging, cells were incubated with 0.5 nM JFX-554 for 15 min in phenol-red-free differentiation medium and washed once before acquisition.

#### Single-molecule imaging

Single-molecule imaging was performed on a Nikon Ti2 inverted microscope equipped with an iLas2 ring-TIRF illumination system and operated in highly inclined illumination (HILO) mode. Images were acquired using a 561-nm excitation beam introduced from a single azimuthal direction at an incidence angle of 54°. The laser was operated at 25% of the instrument output. Fluorescence was collected through a 100× oil-immersion objective (NA 1.49; refractive index 1.515; Nikon ApoChromat) and detected using a Photometrics Fusion sCMOS camera (SN 501487) with 1 × 1 binning and a 10-ms exposure. Images were acquired over a 512 × 512-pixel region at an effective pixel size of 0.065 μm/pixel, corresponding to a 33.28 × 33.28 μm field of view. The camera sensor was maintained at −8°C. Imaging used the microscope L100 light path and 88000v2 quad-band filter set with 1× relay magnification and Nikon Perfect Focus engaged. Movies comprised 5,990 +/- 10 frames acquired over 60 s at ∼100 Hz. Samples were maintained at 37°C and 5% CO₂. Images were acquired using NIS-Elements v5.21.00 and saved as ND2 files.

#### Region selection, single-molecule localization, and trajectory generation

ND2 image sequences were loaded into the ROI-selection module of SPTnano (SPTnano.ROIselector; https://github.com/Michael-shannon/SPTnano; Zenodo link to be added). Regions of interest of 10 μm × 10 μm were selected from each movie and exported as TIFF image stacks which served as input for single particle tracking. Single particles were localized and linked into trajectories using Slimfast (Wong et al. 2025) in MATLAB 2018b. Box size of 22 px and 2 deflation loops were used during localization and tracking. Manual tracking of representative data was used as a reference for comparison of Slimfast with alternative tracking tools, including TrackPy and TrackMate. Slimfast produced the lowest observed tracking error and was therefore used for subsequent analyses.

#### Trajectory preprocessing and motion-feature extraction

Slimfast MATLAB output trajectories were converted to Polars dataframes containing particle identity, frame number, time, and calibrated x- and y-coordinates in micrometers using the SPTnano ParticleMetrics module. Instantaneous and window-level features were then calculated using an HPC workflow in which trajectories were grouped by unique particle identifier, processed in parallel across CPU cores using joblib, and recombined into feature tables.Frame-to-frame displacements were used to calculate instantaneous segment length, speed, direction of motion, acceleration, jerk, curvature, instantaneous diffusion coefficient, and x-and y-velocity components. Undefined values arising from the first frame of a trajectory or from zero-time intervals were set to zero.Local behavioural features were calculated using fixed 60-frame windows with a 30-frame overlap, corresponding to 600-ms windows sampled every 300 ms at 100 Hz. Each window was assigned a unique identifier and analysed independently, allowing transient motion states to be characterized without averaging across complete trajectories. Window-level features included mean speed, self-intersection count, convex hull area, cumulative and net displacement, directional entropy, directional autocorrelation, and turning-angle variance, skewness and kurtosis. Turning angles were normalized to −π to π. Von Mises distributions were fitted to turning angles and absolute directions to estimate directional concentration parameters. Step-length distributions were characterized by their mean, standard deviation, skewness and kurtosis. Additional descriptors included speed variability, instantaneous diffusivity variability, pausing fraction, and the power-spectral-density slope of speed. Diffusive behaviour was quantified using mean-squared displacement (MSD) analysis. For each trajectory or trajectory window, MSD was calculated across lag times and fitted to the anomalous diffusion model MSD(τ) = 4Dτ^α, where D is the diffusion coefficient, τ is the lag time and α is the anomalous exponent. Fits were evaluated in log-log space. Motion was classified as subdiffusive, approximately normal or superdiffusive according to whether α was below, near or above 1, respectively, using a tolerance of 0.1. Poor fits were retained and flagged unless explicitly excluded from subsequent analyses.

#### Definition of behavioural states and pseudo-ground-truth training labels

To identify informative descriptors of molecular behaviour, principal component analysis (PCA) was performed on the extracted motion features. Features contributing most strongly to variance in principal components 1–3 were shortlisted. Pairwise Pearson correlation was subsequently used to identify strongly redundant features within this set. This procedure yielded a reduced feature set comprising mean speed, cumulative displacement, self-intersection count, and the anomalous exponent derived from MSD analysis. Trajectory windows were first manually curated by visual inspection and assigned to four interpretable motion categories: bound, confined, exploratory, and transport-like. These are operational classifications of molecular motion rather than direct measurements of biochemical interactions. Bound trajectories exhibit highly localized movement, consistent with stable molecular engagement, but do not establish binding to a specific partner. Confined trajectories remain within spatially restricted domains, while exploratory trajectories sample larger areas with fewer spatial constraints. Transport-like trajectories exhibit directional displacement consistent with processive movement, although this classification does not independently establish active motor-driven transport. The reduced feature set was then used to construct a hierarchical gating scheme for quantitative assignment of high-confidence examples and to provide a small pseudo-ground truth for training. The first gate used mean speed and self-intersection count to separate major regions of trajectory behaviour. Bound trajectories displayed low speed and high self-intersection counts, whereas exploratory trajectories displayed high speed and low self-intersection counts. Trajectories occupying the relatively low-speed, low-self-intersection region of feature space contained both confined and transport-like behaviours, which were further separated using cumulative displacement and anomalous exponent. Here, low speed refers to the relative position of trajectories within the gated dataset, rather than an absolute definition of transport speed. Only curated high-certainty windows assigned to bound, confined, exploratory, or transport states were used as pseudo-ground-truth labels for supervised transformer training. Unknown or ungated windows were excluded from the supervised classification loss.

#### Raw-motion transformer training and behavioural classification

A compact transformer classifier was trained to assign behavioural states directly from local trajectory dynamics. Each input example corresponded to one trajectory-window identifier and contained up to 60 sequential motion positions.The transformer did not receive the scalar summary features used for manual gating, including mean speed, self-intersection count, cumulative displacement, or anomalous exponent. Instead, trajectories were represented by three local motion channels derived from particle coordinates: frame-to-frame displacement in x, frame-to-frame displacement in y, and directional change as frame-to-frame Δθ. Sequences containing fewer than 60 valid positions were padded, with padding masks supplied to the transformer such that padded positions were excluded during self-attention. Each three-dimensional raw-motion input vector was projected into a 16-dimensional learned embedding. A learnable classification (CLS) token was prepended to the sequence. The resulting sequence was processed by a transformer encoder comprising three encoder layers with four self-attention heads per layer. Each encoder layer contained a feed-forward network that projected the 16-dimensional token representation to 256 dimensions, followed by ReLU activation and projection back to 16 dimensions. The final CLS embedding provides a window-level latent representation of trajectory motion. An offline augmentation-stability screen was performed to identify trajectory perturbations suitable for contrastive transformer training while preserving the behavioural class identity of curated pseudo-ground-truth trajectories. Complete 60-frame trajectory windows were sampled from the gated window table and reconstructed from linked x-y coordinates. For each candidate augmentation, an augmented coordinate path was generated, motion features were recalculated using the same functions used for behavioural gating, and the hierarchical gate geometry was reapplied. Three components of the augmentation procedure were evaluated. First, shuffle severity was varied by dividing each trajectory into increasing numbers of contiguous segments and randomly reordering the segments without applying scaling or angular perturbation. Second, the shuffle was fixed to three segments and the magnitude of bidirectional per-segment displacement scaling was varied. Third, the selected three-segment shuffle and 10% to 50% bidirectional scaling range were retained while per-step angular perturbation was progressively increased.For each augmented window, behavioural-state retention was quantified by determining whether its re-gated behavioural assignment matched its original curated pseudo-ground-truth label. Increasing shuffle severity, scaling magnitude, and angular perturbation progressively increased movement across the manually defined behavioural boundaries. Conservative shuffle-scale-angle parameters produced visible geometric variation while preserving a greater proportion of original behavioural assignments than stronger angular perturbations. Based on this screen, transformer pretraining used a conservative augmentation in which trajectories were divided into contiguous segments, segments were randomly reordered, segment displacements were bidirectionally scaled, and mild per-step angular perturbation was applied. The selected augmentation transform was applied only to valid, unpadded frames: each trajectory was partitioned into contiguous 10-frame segments, each segment was independently rescaled by a factor drawn uniformly from either 0.50–0.90 or 1.10–1.50, the segments were randomly reordered, and each step heading was rotated by 1–10° with a random sign.

Training was then performed in two stages. First, the transformer encoder was pretrained using a contrastive InfoNCE objective. For each trajectory window, an augmented view was generated using the shuffle-scale-angle procedure described above. The original and augmented versions of the same trajectory were treated as a positive pair, whereas other trajectories within the batch were treated as negatives. Negative examples originating from the same unique particle trajectory were excluded. Contrastive pretraining therefore encouraged the encoder to learn trajectory representations that were robust to class-preserving perturbations of local trajectory geometry. The pretrained encoder was subsequently fine-tuned using the curated pseudo-ground-truth behavioural labels. A multilayer perceptron classification head was applied to the CLS embedding and comprised a linear projection from 16 to 32 dimensions, ReLU activation, dropout, and a final linear layer generating four logits corresponding to bound, confined, transport, and exploratory behaviour. Cross-entropy loss was backpropagated through both the classification head and transformer encoder. Validation cross-entropy was calculated on held-out validation data, and the checkpoint with the lowest validation loss was selected for inference. For the final model, validation loss reached its minimum at epoch 98, and this checkpoint was used for subsequent analyses. Initial model-development experiments used 75% of the dataset for training. To determine whether a smaller labelled training population could provide equivalent performance while retaining a larger independent test set, reduced training-set sizes were subsequently evaluated. The final model used a 20% training, 10% validation, and 70% held-out test partition. This reduced-training configuration produced classification accuracy, precision, and confusion-matrix performance comparable with models trained using the larger training set and was therefore used for the analyses presented here. Supervised fine-tuning used only curated pseudo-ground-truth windows within the 20% training partition, corresponding to approximately 5% of the training data. The validation partition was used for checkpoint selection. The 70% held-out test partition was excluded from model training and checkpoint selection and was used only for final inference and evaluation. To visualize the organization of learned motion representations, CLS embeddings from held-out trajectory windows were projected into two dimensions using PHATE with n_components = 2, knn = 5, and decay = 40. Windows were coloured according to their highest-probability behavioural-state prediction. For held-out evaluation of agreement with the feature-based pseudo-ground truth, a random 10% subset of test trajectory windows was labelled using the same hierarchical gating procedure used to generate curated training examples. These independently gated labels were compared with the dominant transformer-predicted state using a confusion matrix. Class-wise agreement was calculated as the proportion of windows within each gated reference class that were assigned to the corresponding transformer-predicted class.

#### Behavioural-state probability and intermediate-state analysis

At inference, the trained classifier generated four logits for each trajectory window. Softmax transformation converted these values into probabilities of membership in the bound, confined, transport, and exploratory states, with probabilities summing to 1 for each window. For each window, probabilities were ranked to identify the most likely class, *P*_top1_, the second-most likely class, *P*_top2_, and their probability margin:

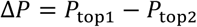

Final behavioural labels were assigned using post hoc probability rules rather than additional learned model parameters. High-confidence single-state assignments were defined using a minimum top-class probability *P*_top1_ 0.85. Boundary-like two-state assignments were defined where substantial probability was distributed across the two highest-scoring states, using a top-two probability margin below 0.4. Windows satisfying neither criterion were retained as unresolved low-confidence predictions. Prediction ambiguity was compared with the manually defined motion-feature space by mapping probability margin onto mean speed versus self-intersection count and cumulative displacement versus anomalous exponent. Representative two-state trajectories and their frequencies across HTT, kinesin, and myosin were subsequently visualized.

#### Long-timescale behavioural-state transition analysis

Longer-timescale behavioural plasticity was quantified from consecutive transformer-defined behavioural-state predictions along individual trajectories. Transformer predictions were generated from 60-frame (600-ms) trajectory windows sampled every 30 frames (300 ms). Consecutive predictions were grouped into sequences of 10 or 20 windows to compare behavioural-state switching over longer temporal scales. Only complete sequences were retained. Each window carried a discrete behavioural label (bound, confined, transport, or exploratory). For each genotype and each superwindow length, conditional transition probabilities *P*(*jj* ∣ *i*) were calculated from successive labels within individual superwindows. State occupancy was the fraction of section assignments in each state within that genotype’s superwindow population. Analyses at 3 s and 6 s were reported separately. To compare switching with the 20H20S control, off-diagonal transition probabilities were expressed as a stabilized percentage change

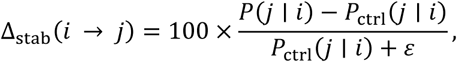

with *εε* = 0.1. The stabilizing term reduced inflation of relative changes for rare transitions while preserving their direction and approximate magnitude. Absolute *P*(*jj* ∣ *i*) values were retained alongside the relative values.

### QUANTIFICATION AND STATISTICAL ANALYSIS

Feature values were first reduced to one median value per cell, with each imaging filename representing one independently analysed cell. Group differences in cell-level medians were assessed using the Kruskal– Wallis test. Where more than two groups were compared, pairwise post hoc comparisons were performed using Dunn’s test with Bonferroni correction. Statistical significance was defined as p < 0.05. Tests were implemented in SciPy. The number of cells and independent experiments, statistical test used, and representation of central tendency and dispersion are specified in the corresponding figure legends.

## References

Andrew, Susan E., Y. Paul Goldberg, Berry Kremer, et al. 1993. “The Relationship between Trinucleotide (CAG) Repeat Length and Clinical Features of Huntington’s Disease.” Nature Genetics 4 (4): 398–403. 10.1038/ng0893-398.

Arrasate, Montserrat, and Steven Finkbeiner. 2012. “Protein Aggregates in Huntington’s Disease.” *Experimental Neurology*, Special issue: Mechanisms of protein misfolding in stressors, vol. 238 (1): 1–11. 10.1016/j.expneurol.2011.12.013.

Aylward, Elizabeth H., Peggy C. Nopoulos, Christopher A. Ross, Douglas R. Langbehn, Ronald K. Pierson, James A. Mills, Hans J. Johnson, Vincent A. Magnotta, Andrew R. Juhl, Jane S. Paulsen, and PREDICT-HD Investigators and Coordinators of Huntington Study Group. 2011. “Longitudinal Change in Regional Brain Volumes in Prodromal Huntington Disease.” Journal of Neurology, Neurosurgery, and Psychiatry 82 (4): 405–10. 10.1136/jnnp.2010.208264.

Aylward, Elizabeth H., Peggy C. Nopoulos, Christopher A. Ross, Douglas R. Langbehn, Ronald K. Pierson, James A. Mills, Hans J. Johnson, Vincent A. Magnotta, Andrew R. Juhl, Jane S. Paulsen, and the PREDICT-HD Investigators and Coordinators of the Huntington Study Group. 2011. “Longitudinal Change in Regional Brain Volumes in Prodromal Huntington Disease.” Research Paper. Journal of Neurology, Neurosurgery & Psychiatry 82 (4): 405–10. 10.1136/jnnp.2010.208264.

Bi, Cheng, Kevin L. Scrudders, Yue Zheng, Maryam Mahmoodi, Shalini T. Low-Nam, and Fang Huang. 2025. “SPTnet: A Deep Learning Framework for End-to-End Single-Particle Tracking and Motion Dynamics Analysis.” Preprint, bioRxiv, February 8. 10.1101/2025.02.04.636521.

Capizzi, Mariacristina, Rémi Carpentier, Eric Denarier, et al. 2022. “Developmental Defects in Huntington’s Disease Show That Axonal Growth and Microtubule Reorganization Require NUMA1.” Neuron 110 (1): 36–50.e5. 10.1016/j.neuron.2021.10.033.

Carpentier, Rémi, Jaesung Kim, Mariacristina Capizzi, et al. 2025. “Structure of the Huntingtin F-Actin Complex Reveals Its Role in Cytoskeleton Organization.” Science Advances 11 (38): eadw4124. 10.1126/sciadv.adw4124.

Caviston, Juliane P., Jennifer L. Ross, Sheila M. Antony, Mariko Tokito, and Erika L. F. Holzbaur. 2007. “Huntingtin Facilitates Dynein/Dynactin-Mediated Vesicle Transport.” Proceedings of the National Academy of Sciences of the United States of America 104 (24): 10045–50. 10.1073/pnas.0610628104.

Chen, Jiji, Zhengjian Zhang, Li Li, et al. 2014. “Single-Molecule Dynamics of Enhanceosome Assembly in Embryonic Stem Cells.” Cell 156 (6): 1274–85. 10.1016/j.cell.2014.01.062.

Dragatsis, I., M. S. Levine, and S. Zeitlin. 2000. “Inactivation of Hdh in the Brain and Testis Results in Progressive Neurodegeneration and Sterility in Mice.” Nature Genetics 26 (3): 300–306. 10.1038/81593.

Duyao, M., C. Ambrose, R. Myers, et al. 1993. “Trinucleotide Repeat Length Instability and Age of Onset in Huntington’s Disease.” Nature Genetics 4 (4): 387–92. 10.1038/ng0893-387.

Duyao, M. P., A. B. Auerbach, A. Ryan, et al. 1995. “Inactivation of the Mouse Huntington’s Disease Gene Homolog Hdh.” *Science* (New York, N.Y.) 269 (5222): 407–10. 10.1126/science.7618107.

Fodale, Valentina, Roberta Pintauro, Manuel Daldin, Maria Carolina Spiezia, Douglas Macdonald, and Alberto Bresciani. 2022. “Quantifying Huntingtin Protein in Human Cerebrospinal Fluid Using a Novel Polyglutamine Length-Independent Assay.” Journal of Huntington’s Disease 11 (3): 291–305. 10.3233/JHD-220527.

Galgoczi, Szilvia, Albert Ruzo, Christian Markopoulos, et al. 2021. “Huntingtin CAG Expansion Impairs Germ Layer Patterning in Synthetic Human 2D Gastruloids through Polarity Defects.” Development (Cambridge, England) 148 (19): dev199513. 10.1242/dev.199513.

Gauthier, Laurent R., Bénédicte C. Charrin, Maria Borrell-Pagès, et al. 2004. “Huntingtin Controls Neurotrophic Support and Survival of Neurons by Enhancing BDNF Vesicular Transport along Microtubules.” Cell 118 (1): 127–38. 10.1016/j.cell.2004.06.018.

Handsaker, Robert E., Seva Kashin, Nora M. Reed, et al. 2025. “Long Somatic DNA-Repeat Expansion Drives Neurodegeneration in Huntington Disease.” Cell 188 (3): 623–639.e19. 10.1016/j.cell.2024.11.038.

Haremaki, Tomomi, Alessia Deglincerti, and Ali H. Brivanlou. 2015. “Huntingtin Is Required for Ciliogenesis and Neurogenesis during Early *Xenopus* Development.” *Developmental Biology*, Modeling Human Development and Disease in Xenopus, vol. 408 (2): 305–15. 10.1016/j.ydbio.2015.07.013.

Haremaki, Tomomi, Jakob J. Metzger, Tiago Rito, M. Zeeshan Ozair, Fred Etoc, and Ali H. Brivanlou. 2019. “Self-Organizing Neuruloids Model Developmental Aspects of Huntington’s Disease in the Ectodermal Compartment.” Nature Biotechnology 37 (10): 1198–208. 10.1038/s41587-019-0237-5.

Kæstel-Hansen, Jacob, Marilina de Sautu, Anand Saminathan, et al. 2025. “Deep Learning-Assisted Analysis of Single-Particle Tracking for Automated Correlation between Diffusion and Function.” Nature Methods 22 (5): 1091–100. 10.1038/s41592-025-02665-8.

Kay, Chris, Jennifer A. Collins, Zosia Miedzybrodzka, et al. 2016. “Huntington Disease Reduced Penetrance Alleles Occur at High Frequency in the General Population.” Neurology 87 (3): 282–88. 10.1212/WNL.0000000000002858.

Kowalek, Patrycja, Hanna Loch-Olszewska, and Janusz Szwabiński. 2019. “Classification of Diffusion Modes in Single-Particle Tracking Data: Feature-Based versus Deep-Learning Approach.” Physical Review E 100 (3): 032410. 10.1103/PhysRevE.100.032410.

Kusumi, Akihiro, Chieko Nakada, Ken Ritchie, et al. 2005. “Paradigm Shift of the Plasma Membrane Concept from the Two-Dimensional Continuum Fluid to the Partitioned Fluid: High-Speed Single-Molecule Tracking of Membrane Molecules.” Annual Review of Biophysics 34: 351–78. 10.1146/annurev.biophys.34.040204.144637.

Laundos, Tiago L., Shu Li, Eric Cheang, Riccardo De Santis, Francesco M. Piccolo, and Ali H. Brivanlou. 2023. “Huntingtin CAG-Expansion Mutation Results in a Dominant Negative Effect.” Frontiers in Cell and Developmental Biology 11 (September). 10.3389/fcell.2023.1252521.

Liu, Zhe, Luke D. Lavis, and Eric Betzig. 2015. “Imaging Live-Cell Dynamics and Structure at the Single-Molecule Level.” Molecular Cell 58 (4): 644–59. 10.1016/j.molcel.2015.02.033.

MacDonald, Marcy E., Christine M. Ambrose, Mabel P. Duyao, et al. 1993. “A Novel Gene Containing a Trinucleotide Repeat That Is Expanded and Unstable on Huntington’s Disease Chromosomes.” Cell 72 (6): 971–83. 10.1016/0092-8674(93)90585-E.

Mätlik, Kert, Matthew Baffuto, Laura Kus, et al. 2024. “Cell-Type-Specific CAG Repeat Expansions and Toxicity of Mutant Huntingtin in Human Striatum and Cerebellum.” Nature Genetics 56 (3): 383–94. 10.1038/s41588-024-01653-6.

Molina-Calavita, Maria, Monia Barnat, Salah Elias, Esther Aparicio, Matthieu Piel, and Sandrine Humbert. 2014. “Mutant Huntingtin Affects Cortical Progenitor Cell Division and Development of the Mouse Neocortex.” Brief Communications. Journal of Neuroscience 34 (30): 10034–40. 10.1523/JNEUROSCI.0715-14.2014.

Piccolo, Francesco M., Nathaniel R. Kastan, Tomomi Haremaki, et al. 2022. “Role of YAP in Early Ectodermal Specification and a Huntington’s Disease Model of Human Neurulation.” eLife 11 (April): e73075. 10.7554/eLife.73075.

Piccolo, Francesco M., Zhe Liu, Peng Dong, et al. 2019. “MeCP2 Nuclear Dynamics in Live Neurons Results from Low and High Affinity Chromatin Interactions.” eLife 8 (December): e51449. 10.7554/eLife.51449.

Pinholt, Henrik D., Søren S. R. Bohr, Josephine F. Iversen, Wouter Boomsma, and Nikos S. Hatzakis. 2021. “Single-Particle Diffusional Fingerprinting: A Machine-Learning Framework for Quantitative Analysis of Heterogeneous Diffusion.” Proceedings of the National Academy of Sciences 118 (31): e2104624118. 10.1073/pnas.2104624118.

Quintanilla, Melissa A., John A. Hammer, and Jordan R. Beach. 2023. “Non-Muscle Myosin 2 at a Glance.” Journal of Cell Science 136 (5): jcs260890. 10.1242/jcs.260890.

Reiner, A., R. L. Albin, K. D. Anderson, C. J. D’Amato, J. B. Penney, and A. B. Young. 1988. “Differential Loss of Striatal Projection Neurons in Huntington Disease.” Proceedings of the National Academy of Sciences of the United States of America 85 (15): 5733–37. 10.1073/pnas.85.15.5733.

Ross, Christopher A., and Sarah J. Tabrizi. 2011. “Huntington’s Disease: From Molecular Pathogenesis to Clinical Treatment.” The Lancet. Neurology 10 (1): 83–98. 10.1016/S1474-4422(10)70245-3.

Saudou, Frédéric, and Sandrine Humbert. 2016. “The Biology of Huntingtin.” Neuron 89 (5): 910–26. 10.1016/j.neuron.2016.02.003.

Shannon, Michael J., Shira E. Eisman, Alan R. Lowe, Tyler F. W. Sloan, and Emily M. Mace. 2024. “cellPLATO – an Unsupervised Method for Identifying Cell Behaviour in Heterogeneous Cell Trajectory Data.” Journal of Cell Science 137 (20): jcs261887. 10.1242/jcs.261887.

Shannon, Michael J., Judith Pineau, Juliette Griffié, et al. 2020. “Differential Nanoscale Organisation of LFA-1 Modulates T-Cell Migration.” Research Article. Journal of Cell Science 133 (5). 10.1242/jcs.232991.

Vaswani, Ashish, Noam Shazeer, Niki Parmar, et al. 2017. “Attention Is All You Need.” Advances in Neural Information Processing Systems 30. https://proceedings.neurips.cc/paper_files/paper/2017/hash/3f5ee243547dee91fbd053c1c4a845aa-Abstract.html.

Vega, Anthony R., Spencer A. Freeman, Sergio Grinstein, and Khuloud Jaqaman. 2018. “Multistep Track Segmentation and Motion Classification for Transient Mobility Analysis.” Biophysical Journal 114 (5): 1018–25. 10.1016/j.bpj.2018.01.012.

Verhey, Kristen J., and Ryoma Ohi. 2023. “Causes, Costs and Consequences of Kinesin Motors Communicating through the Microtubule Lattice.” Journal of Cell Science 136 (5): jcs260735. 10.1242/jcs.260735.

Zhang, Yu, Blair R. Leavitt, Jeremy M. van Raamsdonk, et al. 2006. “Huntingtin Inhibits Caspase-3 Activation.” The EMBO Journal 25 (24): 5896–906. 10.1038/sj.emboj.7601445.

