## Supplementary figures for "Single-molecule motion profiling reveals direct and indirect effects of PolyQ expansion on huntingtin dynamics and molecular engagement"

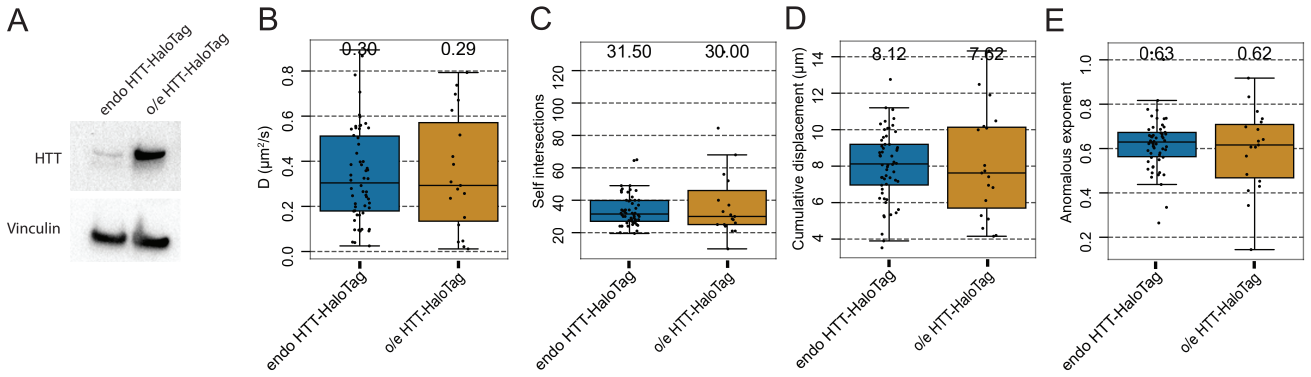


Figure S1: Endogeneously tagged HTT has the same diffusion pattern as overexpressed tagged HTT. A) Western blot of endogeneous HTT-HaloTag. Boxplots showing endogeneous vs overexpressed HTT-HaloTag single particle track measurements of B) diffusion coefficient, C) self-intersections, D) cumulative displacement and E) anomalous exponent. Dots = mean per cell.

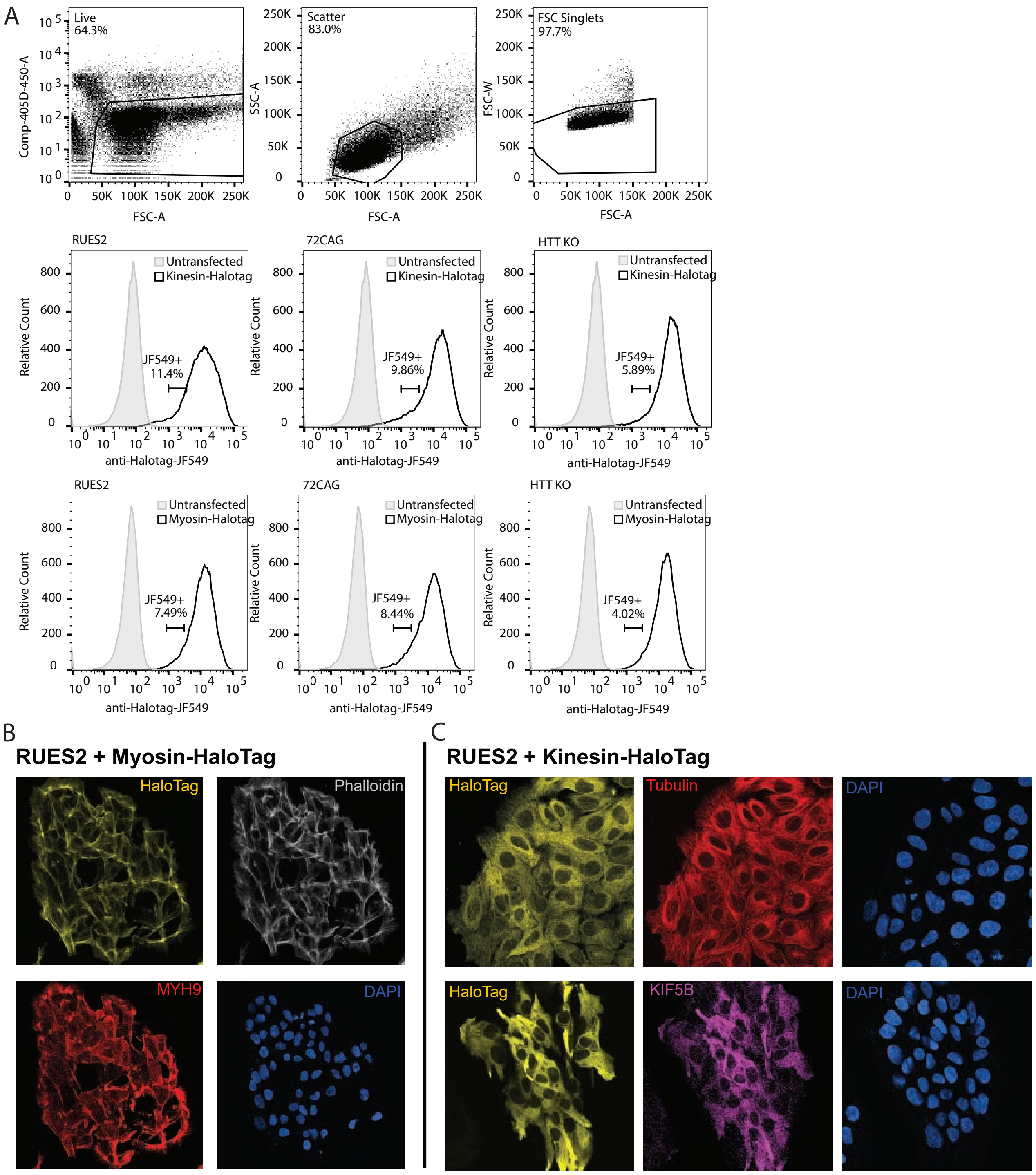


Figure S2: FACS, imaging and biochemical confirmation of kinesin-halotag containing RUES2, 72CAG and HTTKO lines. A) FACS gating strategy for isolating low expression populations of kinesin and myosin halotag lines on RUES2 (WT HTT), 72CAG-HTT and HTT-KO backgrounds. C) Representative confocal images of hESCs stained with HaloTag-ligand, phalloidin, myosin antibody and DAPI. D) Representative confocal images of hESCs stained with HaloTag-ligand, tubulin antibody, and DAPI in the top row; bottom row HaloTag-ligand, Kinesin antibody and DAPI.

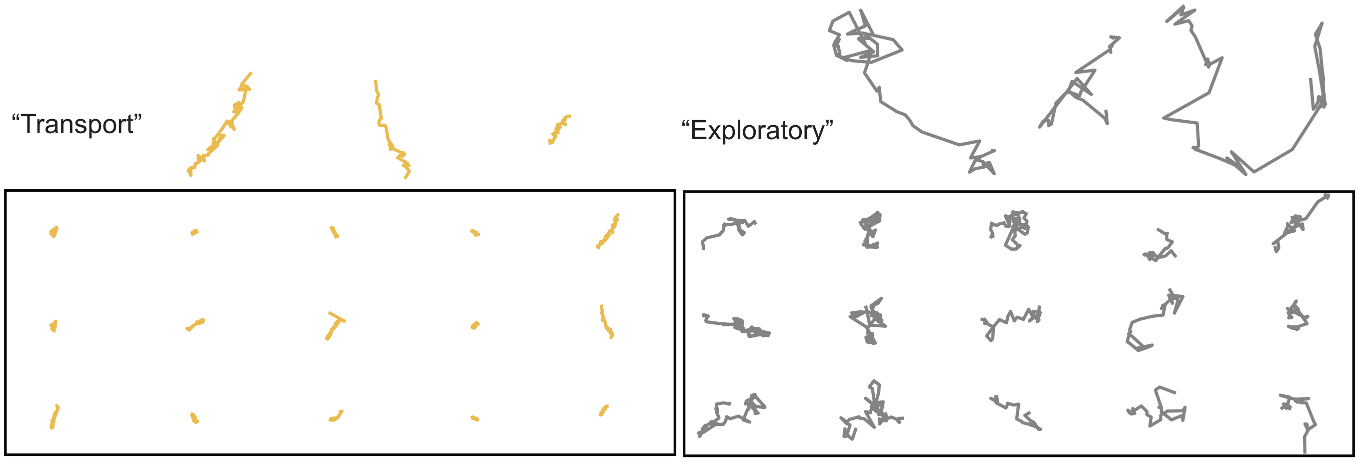


Figure S3: Representative examples of behaviour types manually selected as pseudo-ground truth labels. Each behaviour is represented as a time windowed length of a track: all are 600 ms (60 frames) in length. A) Bound, B) confined, C) transport, D) exploratory.

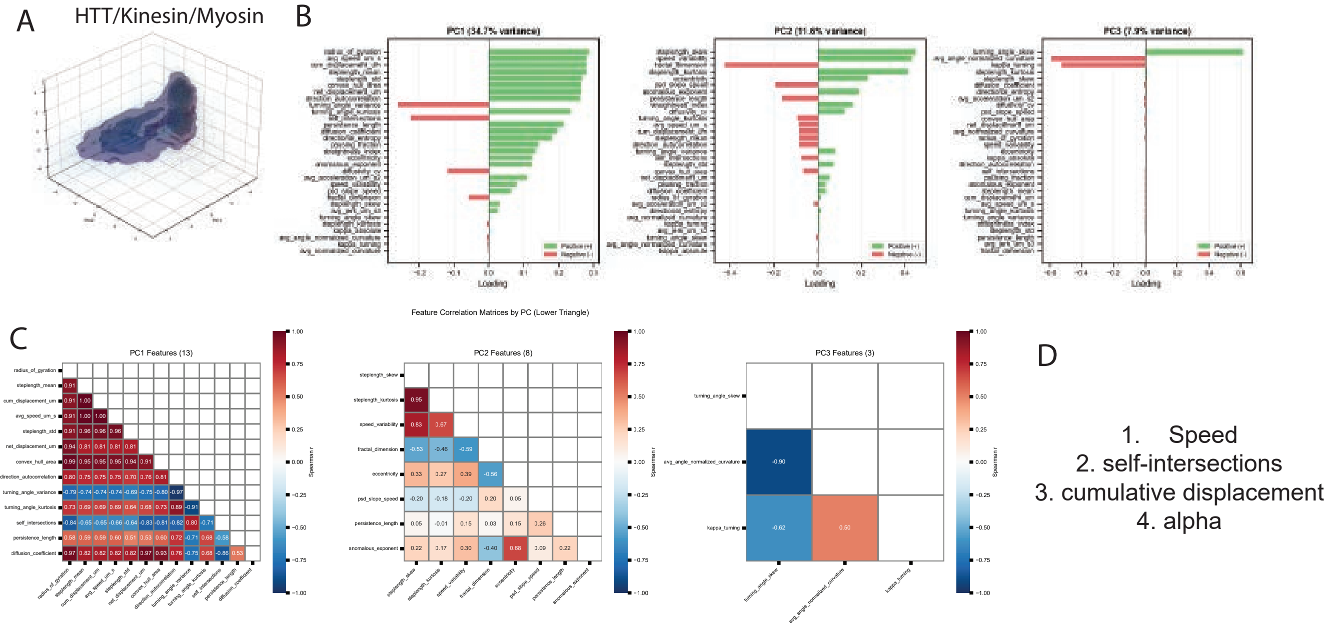


Figure S4: Feature selection for non-redundant description of molecular tracks. A) Features were assessed by PCA, and B) their contribution to the variance of the first 3 PC values was extracted. C) Redundancy was assessed by plotting the correlation between features and selecting those with close to 0 for each PC to create D) a shortlist of high variance, low redundancy interpretable features.

### Augmentation gallery

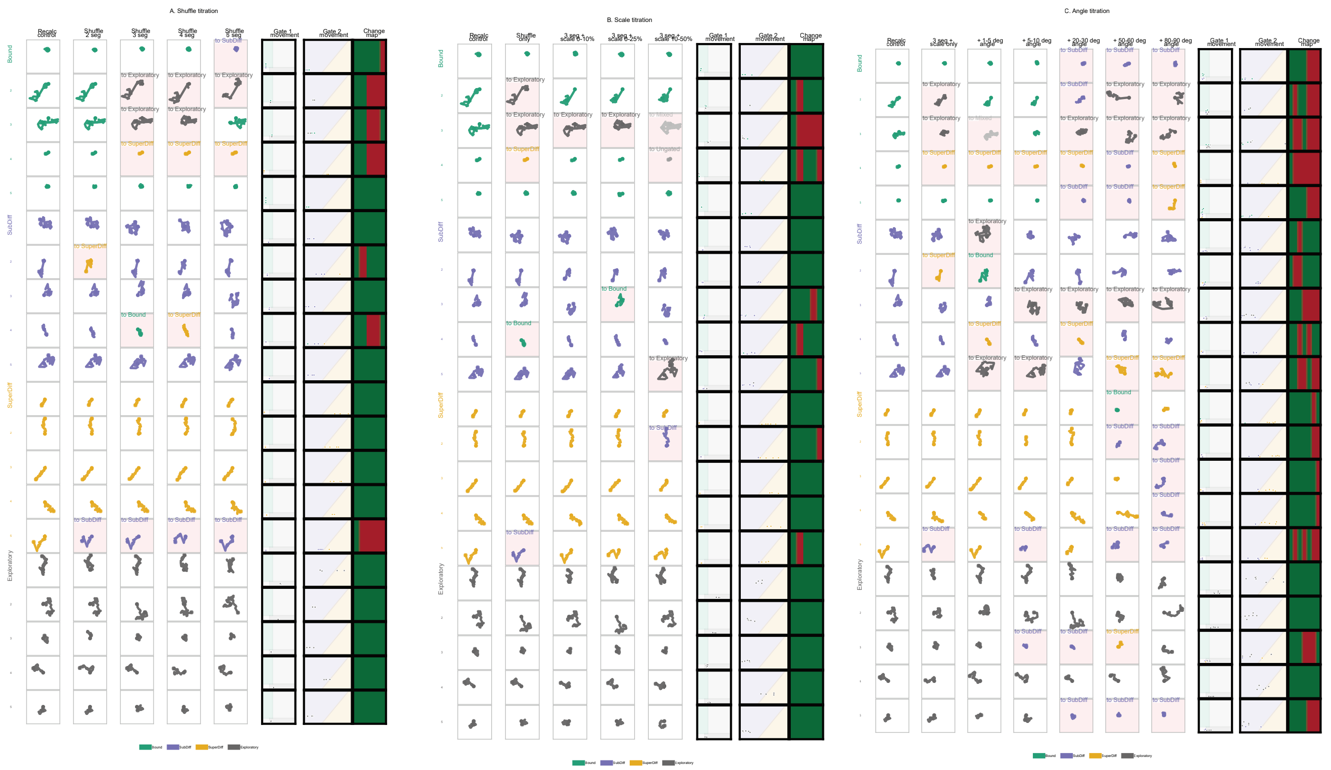


Fig. S5. Offline screen of raw-trajectory augmentation parameters. Complete 60-frame windows were sampled from the curated pseudo-ground-truth behavioural populations. Each row shows one parent trajectory window and each column shows either the recalculated control or an augmented version of the same window. Trajectory color indicates the population assigned after recomputing the gating features and reapplying the hierarchical gate geometry. Pale red backgrounds and the text annotations indicate augmentations that changed category. The final columns show the positions of the recalculated and augmented windows in Gate 1 feature space, Gate 2 feature space, and a binary heatmap of whether category assignment changed. (A) Shuffle titration, varying the number of shuffled trajectory segments while leaving scaling and angular perturbation off. (B) Scale titration, using the selected 3-segment shuffle while increasing bidirectional per-segment scaling. (C) Angle titration, using the selected 3-segment shuffle and 10-50% segment scaling while increasing per-step angular perturbation. (D) Larger-sample summary showing the percentage of sampled windows that changed category under each perturbation, split by population; circled points indicate the mild shuffle-scale-angle settings selected for transformer training.

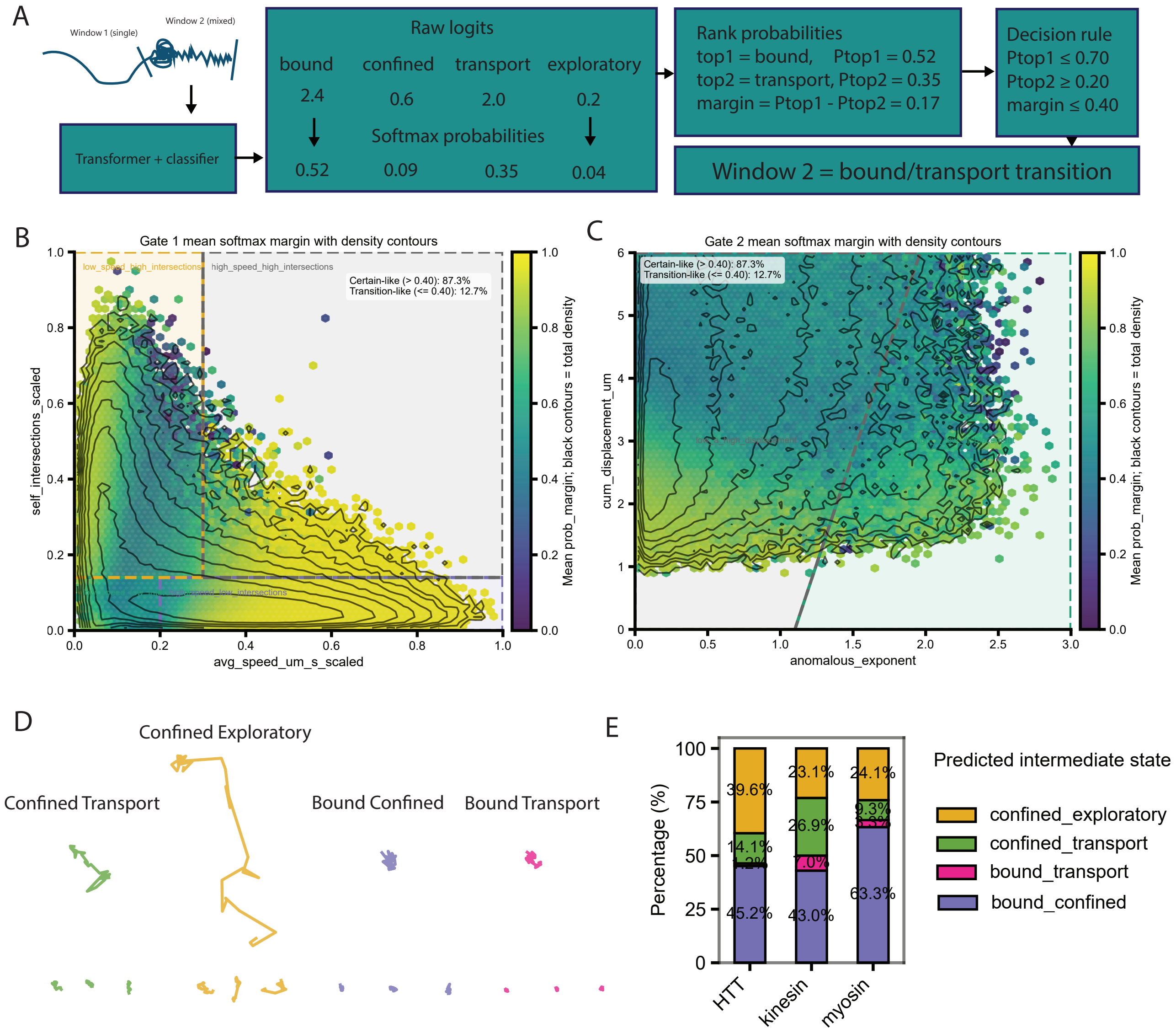


Figure S6: Intermediate states identified using transformer. A) Diagram showing method of extracting probabilities to define mixed states. B) Hex and contour map of speed vs self intersections and C) cumulative displacement vs anomalous exponent, coloured by probabilitiy margin. A high margin indicates single state, a low margin (below threshold 0.4) indicates transition states. D) Transition state track examples. E) Intermediate state distributions in ES cells per molecule.

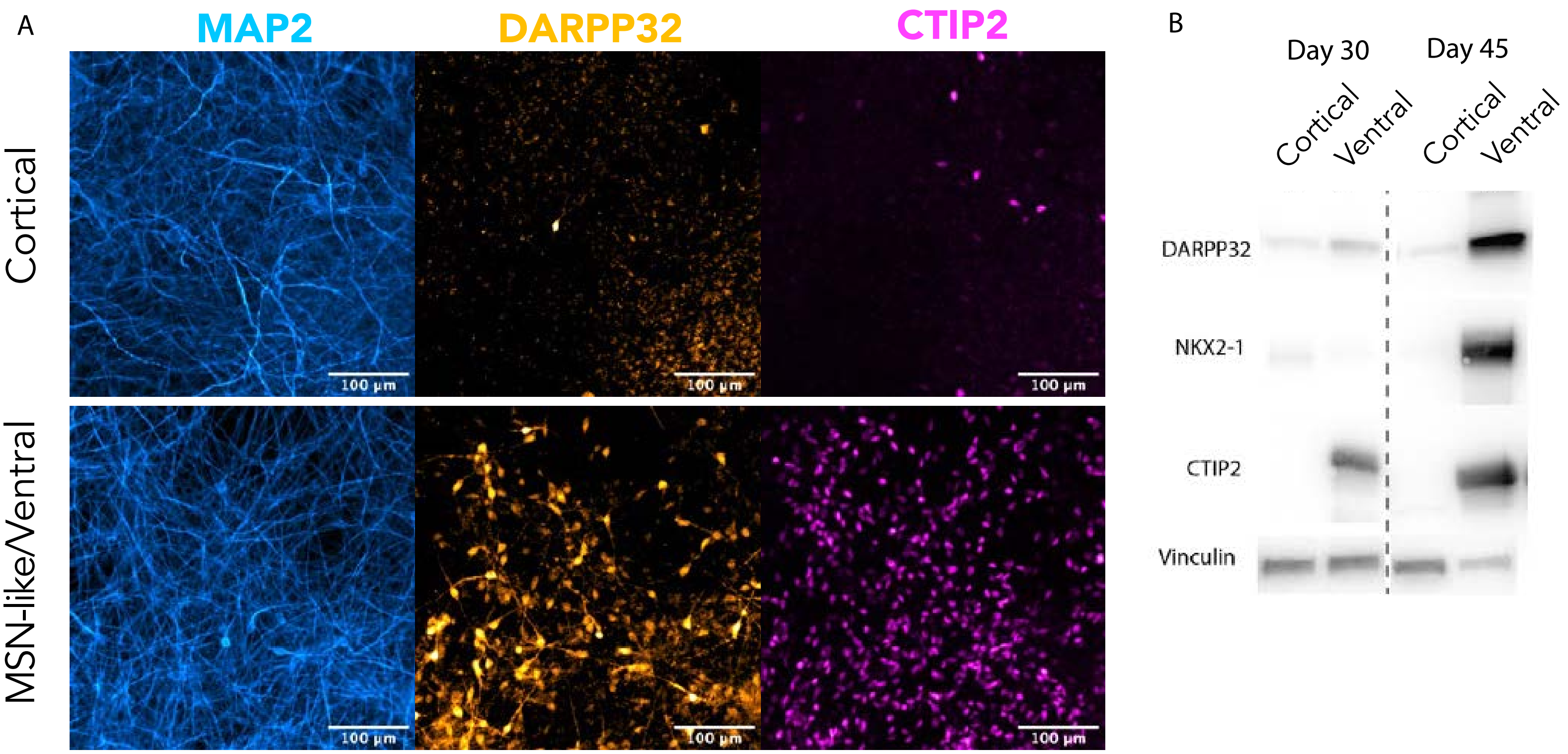


Figure S7: Neurons differentiated from RUES2 cells display markers consistent with Cortical and MSN-like/Ventral neurons. A) Confocal microscopy images of differentiated neurons stained for MAP2, DARPP32 and CTIP2; top row = cortical, bottom row = MSN-like/ventral. B) Western blot of cortical vs ventral neurons at day 30 and day 45 of differentiation showing DARPP32, NKX2-1, CTIP2 and Vinculin.

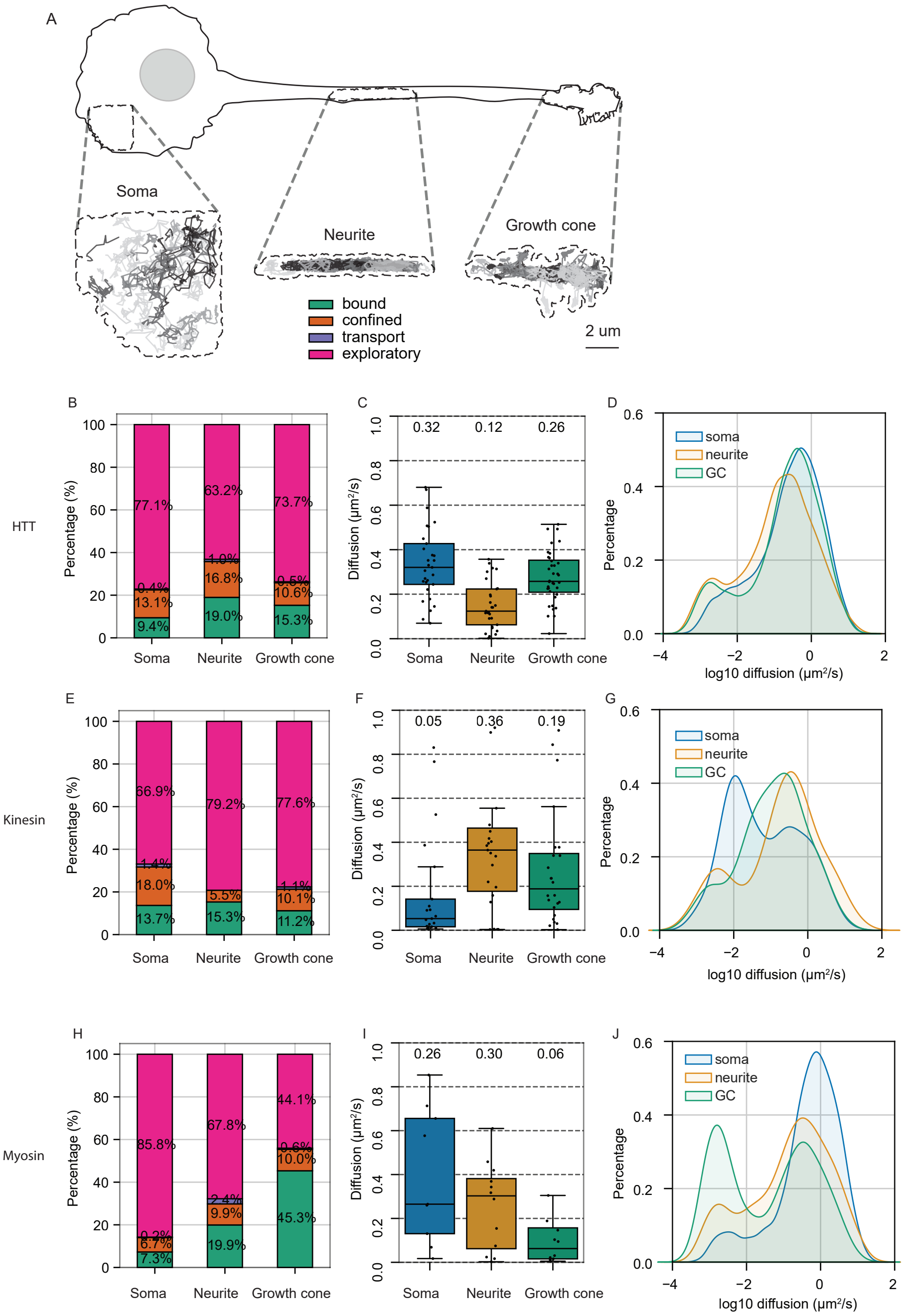


Figure S8: HTT, Kinesin and Myosin have distinct track profiles regionally within cortical neurons. A) HTT was imaged in the soma, neurite and growth cone and a subset of tracks are displayed - contours display the track maxima from zoomed ROIs. B) HTT track fingerprint plot of the percentage proportion of each motion class in the soma, neurite and growth cone. C) Boxplot of HTT diffusion coefficient per region. D) Histogram of HTT log10 diffusion coefficient per region. E, F, G) Same for Kinesin. H, I, J) same for myosin.

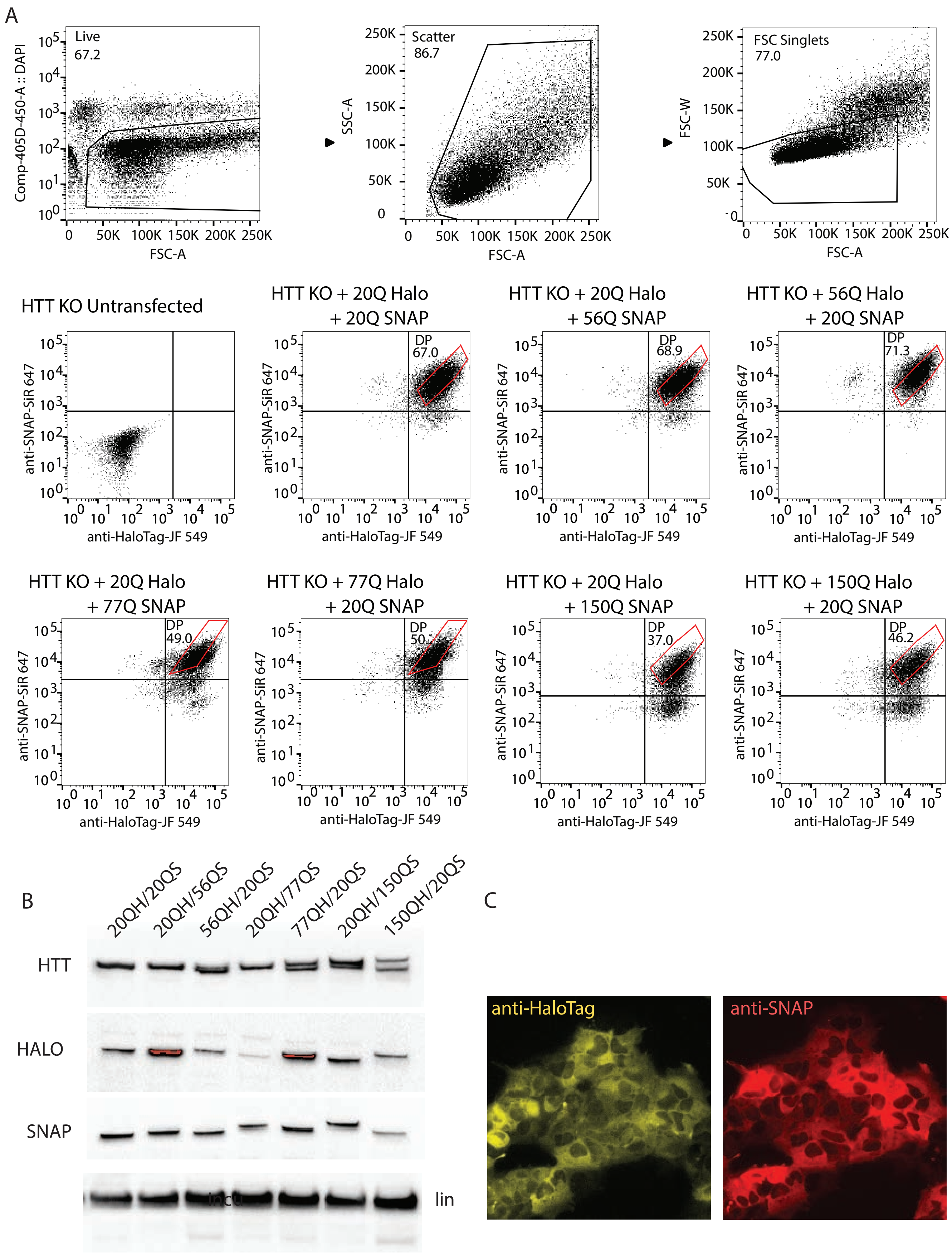


Figure S9: FACS and validation strategy for the HTT tag-swap allelic series. A) gating strategy B) western blot, C) confocal imaging.

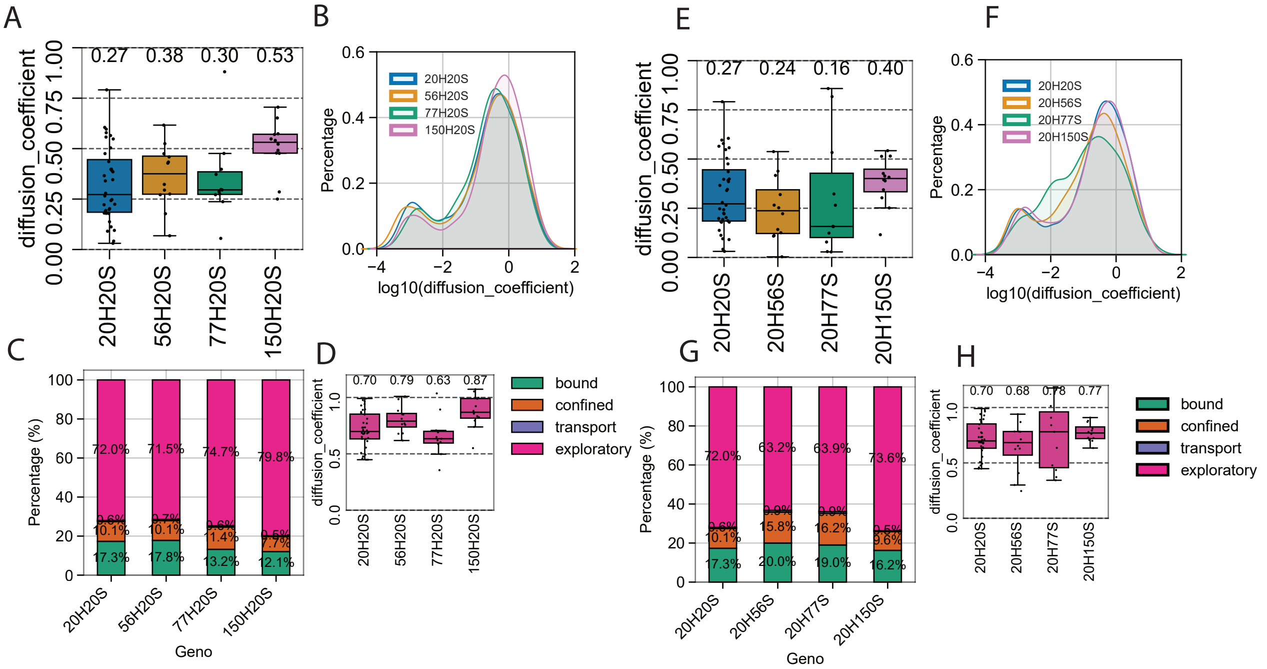


Figure S10: PolyQ-expanded HTT alters HTT dynamics in hESCs. A–D) Diffusion coefficient, log10 diffusion-coefficient distribution, motion-class fingerprint and exploratory-population diffusion coefficient for directly tracked 20Q, 56Q, 77Q and 150Q HTT. E–H) Corresponding analyses of wild-type 20Q HTT co-expressed with 20Q, 56Q, 77Q or 150Q HTT.

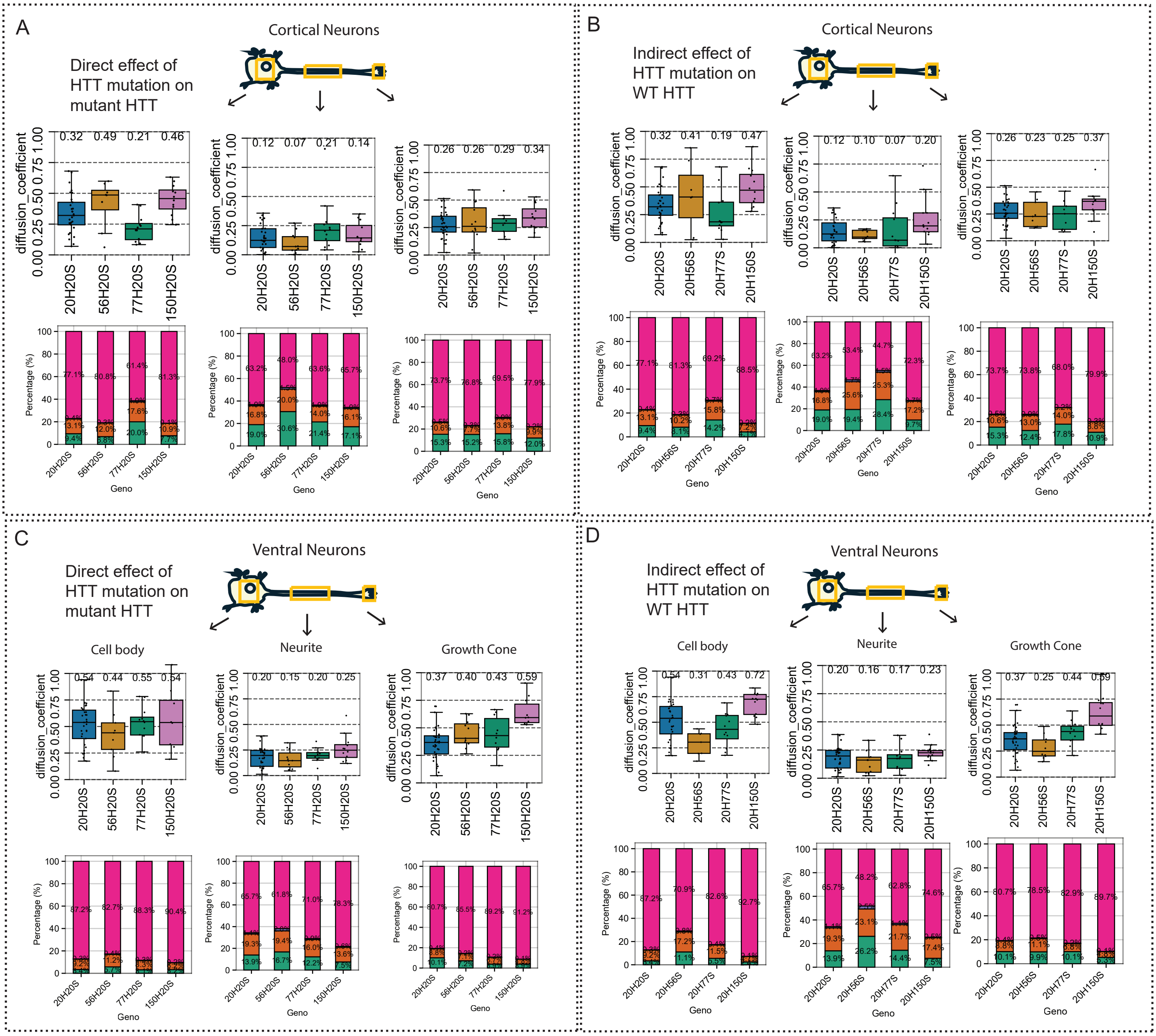


Figure S11: PolyQ-dependent changes in HTT dynamics measured across the cortical and MSN-like soma, neurite and growth cone. Diffusion coefficient of HTT in the soma, neurite and growth cone across the PolyQ allelic series, and corresponding motion-class fingerprints showing the proportions of bound, confined, exploratory and transport-like trajectories. Shown for A) direct effects of mutation in cortical neurons, B) indirect effects of HTT mutation on WT HTT in cortical neurons, C) direct effects of mutation in ventral neurons, and D) indirect effects of mutation in ventral neurons.

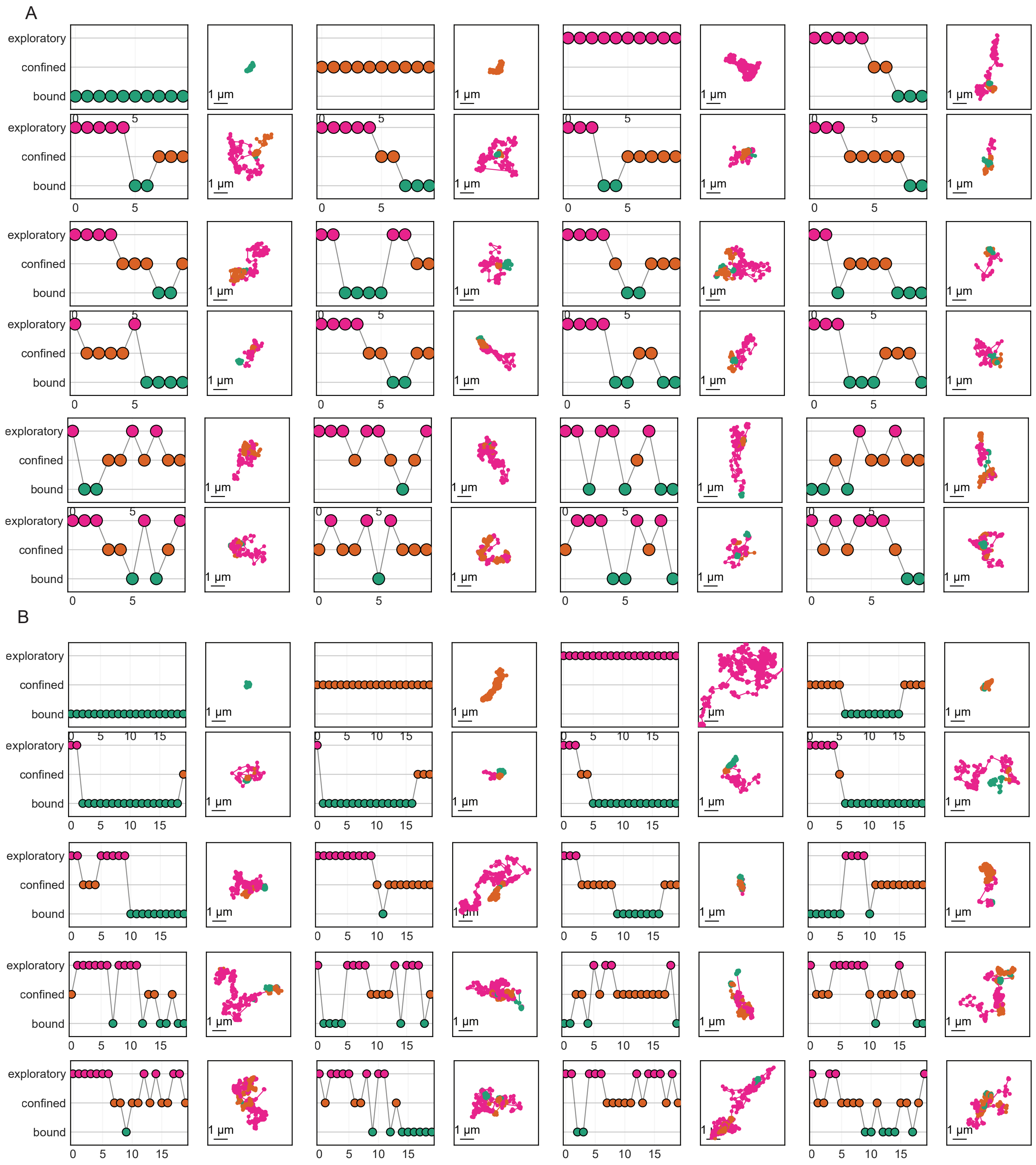


Figure S12: Gallery of track transitions and accompanying tracks coloured by state. Superwindows were made up of 10 (A) or 20 (B) sequential windows. Each window was 300 ms long, composed of 30 frames. Superwindows were therefore 3 or 6 seconds long, composed of 300 or 600 frames.

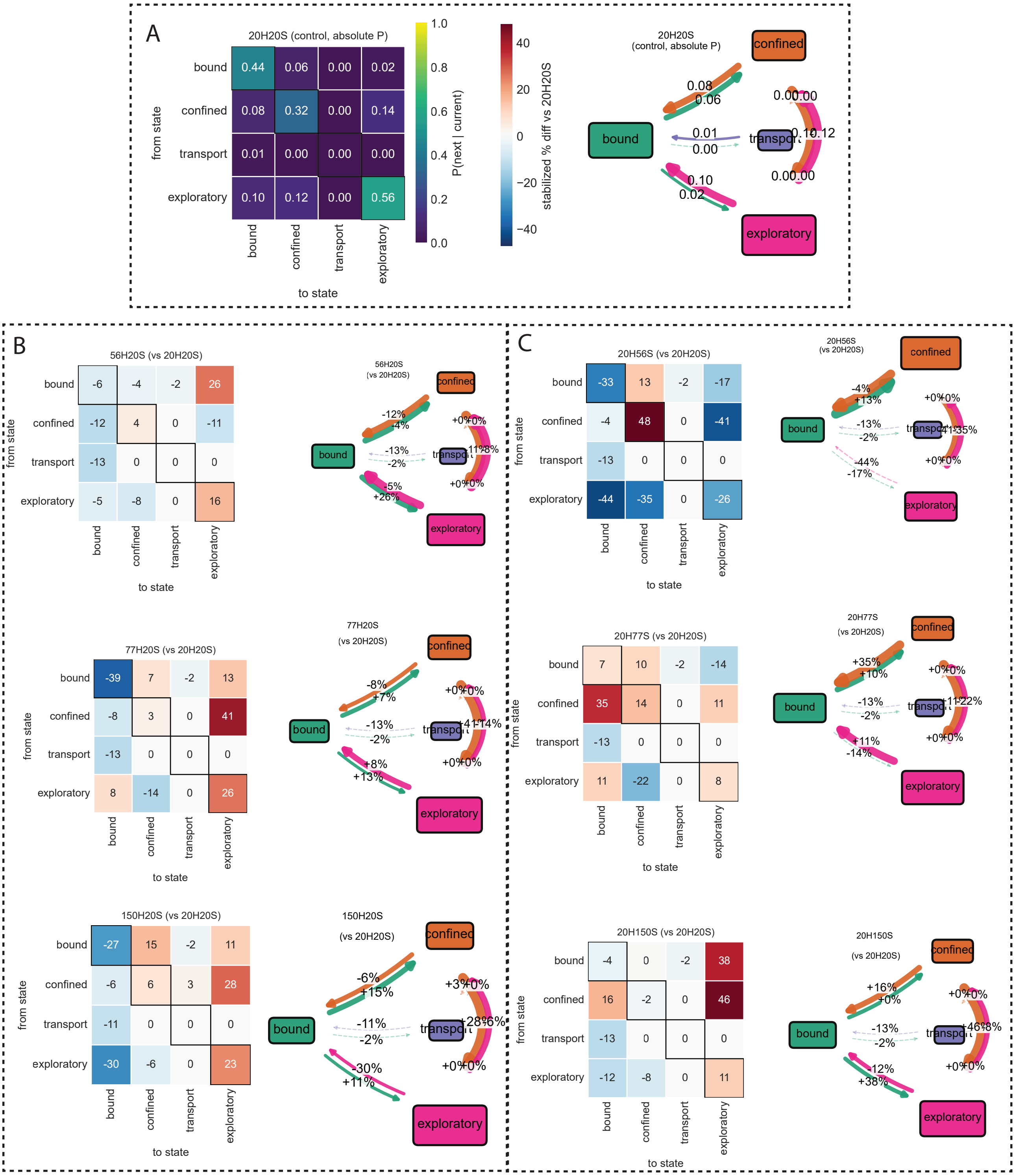


Figure S13: Transition switches in the ventral neuron growth cone. A) Probability heatmap and resultant fan diagram for 20H20S ventral neuron growth cones. B) Percent change in state switch probability in heatmaps and fan diagrams for the direct effect of PolyQ mutation on mutant HTT. C) Same for the indirect effect of PolyQ mutation on wild-type HTT.

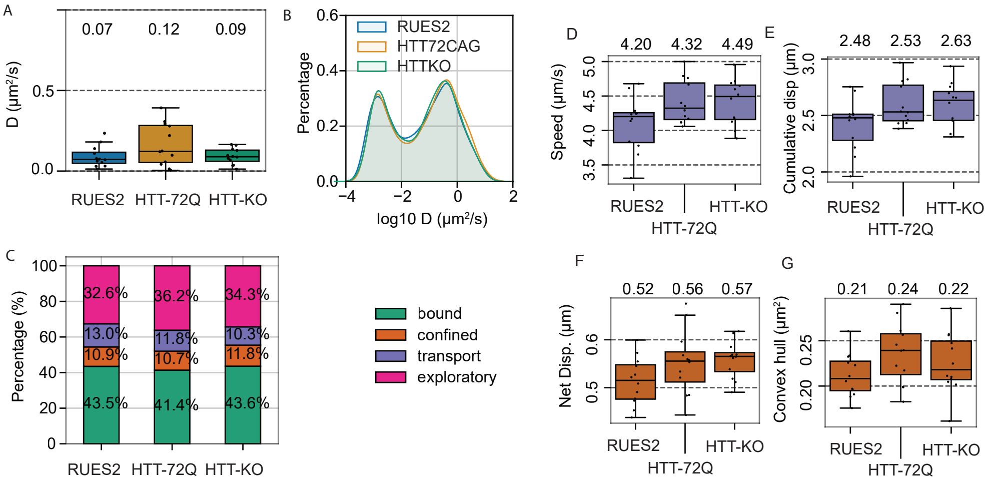


Figure S14: HTT mutation or loss alters kinesin motion states in ES cells. A) Boxplot of kinesin diffusion coefficient in ES cells with HTT-WT, HTT-72Q or HTT-KO backgrounds. B) Histogram of kinesin log10 diffusion coefficient in each HTT background. C) Fingerprint plot showing the percentage of trajectories assigned to each predicted motion class. D) Speed of trajectories assigned to the transport class. E) Net displacement of trajectories assigned to the confined class. F) Convex hull area of trajectories assigned to the confined class.

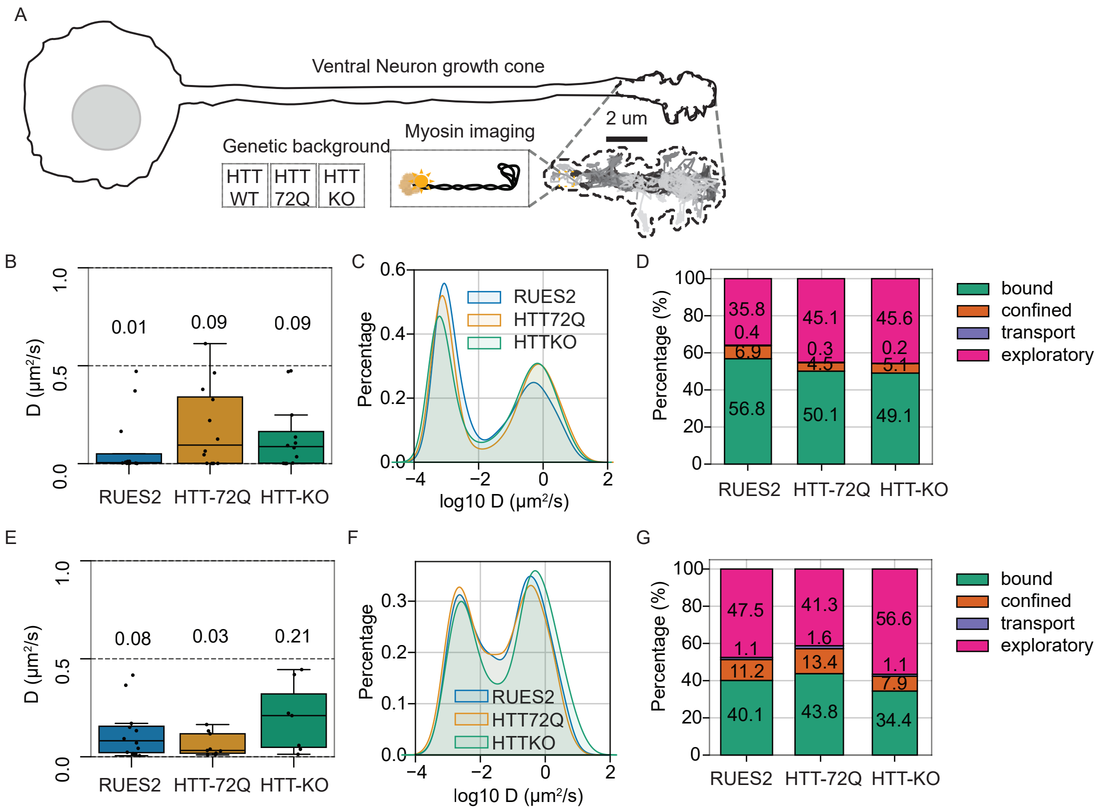


Figure S15: HTT mutation or loss reduces myosin binding in ventral-neuron growth cones and ES cells. A) Diagram of set up B) Boxplot of myosin diffusion coefficient in hESC at pluripotency with HTT-WT, HTT-72Q or HTT-KO backgrounds. C) Log10 diffusion coefficient KDE histogram. D) Fingerprint plot showing the percentage of myosin trajectories assigned to each predicted motion class in ventral-neuron growth cones. E, F and G) same for the growth cones of MSN-like/ventral-neuron.

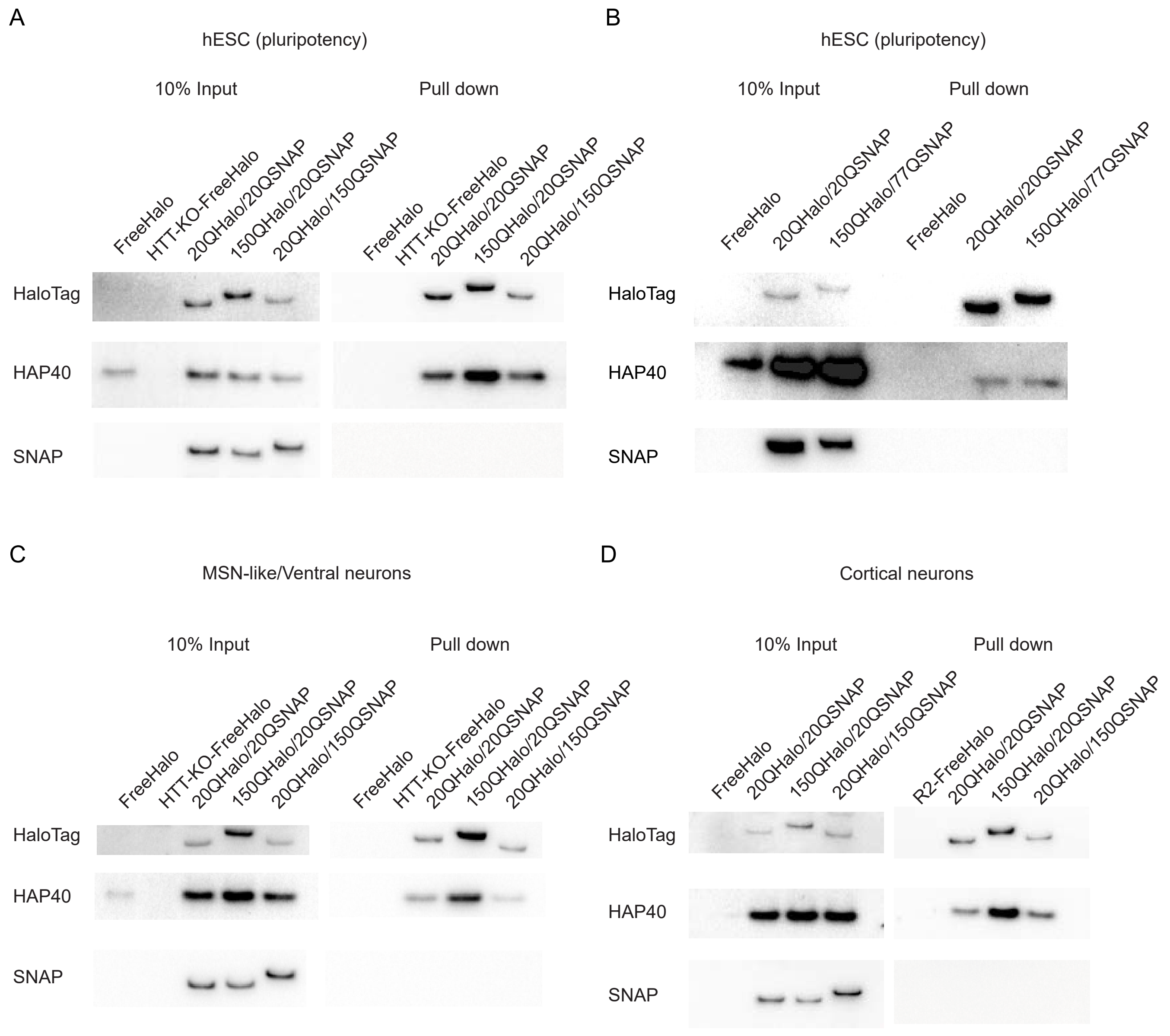


Figure S16: HTT does not dimerize with other HTT proteins. Immunoblots for HaloTag, HAP40 and SNAP in total extract (10% Input) and immunoprecipitation with anti-HaloTag nanobodies (Pull down) in differentially tagged cells. A, C and D) Pull down of HTT-HaloTag in 20QHalo/20SNAP (wtHTT in non-HD), 150QHalo/20QSNAP (mutant HTT), 20QHalo/150QSNAP (wt HTT in HD context) at pluripotency (A), in MSN-like/Ventral (C) and Cortical neurons (D). B) Pull down of HTT-Halo in control cells 20QHalo/20SNAP (wtHTT in non-HD) and in cells expressing two forms of mutant HTT (150 Halo and 77 SNAP) at pluripotency. In each condition, cells expressing HaloTag alone in wt (freeHalo) or in HTT-KO (HTT-KO-FreeHalo) were used as negative control for the pull down. The fact that immunoprecipitation of HTT-HaloTag pulls down HAP40 (the best known HTT interactors) indicate that the pull down was successful. Because no SNAP signal was detectable in the IP, this indicates that HTT does not dimerize in these experimental conditions.
